# Correlative light and electron microscopy identifies circuit motifs underlying processing of noisy visual motion in larval zebrafish

**DOI:** 10.1101/2025.03.14.643363

**Authors:** Jonathan Boulanger-Weill, Florian Kämpf, Gregor F. P. Schuhknecht, Richard L. Schalek, Roberto Garza, Mariela Petkova, Sumit Kumar Vohra, Yuelong Wu, Jay H. Savaliya, Robert Tiller, Kristian J. Herrera, Heike Naumann, Maren Eberle, Simone Rencken, Moritz Stingl, Alina Hebling, Dana Hockling, Katja Slangewal, Zixuan Deng, Ruohong C. Wang, Lauren L. Zhang, Kim N. Kirchberger, Isaac H. Bianco, Daniel Baum, Adrienne L. Fairhall, Filippo Del Bene, Florian Engert, Jeff W. Lichtman, Armin Bahl

## Abstract

To integrate noisy visual motion, vertebrate circuits must transform transient sensory signals into persistent, lateralized activity. The synaptic architecture supporting this computation remains unclear. We combined calcium imaging and correlative electron microscopy (EM) in larval zebrafish with an independent whole-brain EM dataset, reconstructing 1,246 neurons in and connected to the anterior hindbrain, a region previously implicated in the temporal integration of visual evidence. We identified four broad functional and morphological classes interconnected through recurrent excitation, interhemispheric and ipsilateral inhibitory motifs, as well as long-range inputs from rostral visual regions. We then trained bilateral neural networks constrained by the measured neuron-to-neuron connectivity as a fixed anatomical mask, while optimizing unknown physiological weights. These models reproduced population dynamics and predicted responses to previously untested motion stimuli. Shuffling connectivity or selectively removing recurrent connections impaired model performance, whereas removing an equivalent number of random connections across the network had much smaller effects. Our results identify an anatomical network compatible with visual-motion integration and implicate specific recurrent connections in its computational dynamics. Together, our results link functional dynamics with synaptic architecture in a vertebrate sensorimotor circuit and provide a framework for interpreting connectomic datasets through combined functional, anatomical, and computational analyses.

## Introduction

In natural environments, behaviorally relevant signals are often noisy, fragmented, or conflicting. Producing an appropriate action therefore requires several coordinated computations: sensory evidence must be integrated across space and time, competing action representations must be compared, and motor pathways must be engaged only when sufficient evidence is available. A classic approach for studying this process is through random dot motion stimuli, where some dots move coherently while others are randomly redrawn, flickering across the visual field. This stimulus has been used alongside single-neuron and population-level recordings to reveal ramping neural responses, with slopes correlated with coherence levels, indicative of sensory evidence accumulation (Newsome, Britten and Movshon, 1989; Hanks and Summerfield, 2017). Recent findings suggest that this integration is distributed throughout the mammalian brain, involving both cortical and subcortical regions (Brody and Hanks, 2016). However, the detailed architecture, both at the synaptic and whole-brain levels, that transforms noisy sensory input into discrete motor actions in vertebrates remains unclear.

The zebrafish larva (*Danio rerio*) also responds to random dot motion, accumulating visual evidence of self-motion before initiating stabilizing motor commands (Bahl and Engert, 2020; Dragomir, Štih and Portugues, 2020). Its compact brain and genetic amenability make it a powerful model for investigating neural computations using light microscopy, enabling whole-brain functional circuit dissection that is not feasible in larger vertebrates. Previous research has highlighted the anterior hindbrain, where neurons display calcium dynamics indicative of visual evidence accumulation, as a key computational hub for this process (Bahl and Engert, 2020; Dragomir, Štih and Portugues, 2020). Several network predictions have been proposed: hindbrain neurons accumulate visual input from pretectal (Pt) regions by recurrent connectivity of excitatory neurons, generating persistent activity (Yang *et al*., 2023); faster-dynamics neurons establish a decision boundary by preventing premature motor activation (Bahl and Engert, 2020) and motor-command neurons select directionally appropriate reticulospinal populations through excitatory and inhibitory connections (Orger *et al*., 2008; Kubo *et al*., 2014; Naumann *et al*., 2016). Together, these predictions suggest that noisy-motion processing involves integration, supported by the coordinated operation of multiple connectivity motifs.

Volumetric electron microscopy techniques have made it possible to map synaptic connectivity with remarkable resolution in several invertebrate and vertebrate species (Witvliet *et al*., 2021; Svara *et al*., 2022; Turner *et al*., 2022; Dorkenwald *et al*., 2024; Schlegel *et al*., 2024; Shapson-Coe *et al*., 2024). In *Drosophila*, this connectomic data, when combined with complementary methods such as electrophysiology, optogenetics, and behavioral analysis, can refine models of neural networks (Turner-Evans *et al*., 2020; Ammer *et al*., 2023), predict neural dynamics (Lappalainen *et al*., 2024; Shiu *et al*., 2024), and elucidate the mechanistic basis of behavior (Dombrovski *et al*., 2023; Shiu *et al*., 2024). In vertebrates, neurons are far less stereotypic and do not lend themselves as readily to assigning function based on structure. Further, similar functional response types are often scattered across regions, making it difficult to test proposed wiring schemes through sparse connectivity sampling from separate animals. A prime example of such a lack of topographic representation is the zebrafish pretectum, where binocular and monocular neurons are intermingled (Kubo *et al*., 2014; Naumann *et al*., 2016). Nonetheless, recent advances in functional connectomics in this model organism—combining in vivo calcium imaging with subsequent electron microscopy in the same animal—have made it possible to directly link functional neuronal activity with its underlying synaptic connectivity (Wanner and Friedrich, 2020; Friedrich and Wanner, 2021; Vishwanathan *et al*., 2024). This approach offers unprecedented insights into how connectomic motifs support neuronal responses, sensory processing, and ultimately behavior in the vertebrate brain.

Building on previous work in which we had proposed a plausible computational model for processing of noisy visual sensory evidence in the vertebrate brain (Bahl and Engert, 2020), we now use ground-truth synaptic connectivity to investigate the circuit architecture underlying these computations. Specifically, we generated connectivity matrices from two separate EM volumes. The first dataset, a functional correlated EM (FCLEM) volume, was obtained after functionally imaging the anterior hindbrain during random dot motion stimuli. The second one spans the whole brain (WBEM), enabling circuit mapping across broader networks, but lacks functional information. To bridge structure and function, we created a library of photoactivated neurons with known activity profiles and used it to train a classifier capable of predicting broad functional identity from morphology in the anterior hindbrain. Combining these approaches, we identify recurrent excitation, interhemispheric inhibition, and ipsilateral disinhibition as core motifs associated with processing of motion signals. We further investigated whether the reconstructed connectivity provides an anatomical substrate compatible with the observed temporal integration dynamics. Using connectome-constrained recurrent network models, we reproduced the measured population dynamics and predicted responses to novel visual stimuli. Targeted connectivity perturbations further identified recurrent excitation among ipsilateral motion-integrating neurons as particularly important for supporting slow integration within the model. These findings refine our understanding of how noisy sensory signals are transformed into motor outputs and provide a framework for linking synaptic connectivity to circuit dynamics in vertebrate brains.

## Results

### Correlative electron and light microscopy in a neural circuit performing motion integration

To explore the relationship between structure and function in the anterior hindbrain, we first identified neurons based on their functional dynamics using two-photon calcium imaging (**Fig. 1a** and (Bahl and Engert, 2020)). To that end, we used a 7 days post-fertilization (dpf) transgenic larvae expressing a nuclear calcium indicator (Dana *et al*., 2019) under the *elavl3* pan-neuronal promoter and red markers in the main inhibitory neuronal subtype (Satou *et al*., 2013) as well as in vascular endothelial cells (Fujita *et al*., 2011), *Tg(elavl3:H2B-GCaMP7f, gad1b:DsRed, kdrl:mCherryCAAX)*. We recorded neuronal responses evoked by left- and rightward-moving dots of 100% coherence in six optical planes spaced by 12 μm along the dorso-ventral axis in a region spanning from the medial optic tectum to the caudal hindbrain. We performed 2D segmentation of the time-averaged recordings using a convolutional neural network (Weigert *et al*., 2020) to obtain neuron-specific masks (**Methods**). While many diverse neurotransmitter types exist (Higashijima, Mandel and Fetcho, 2004), the transgenic Gad1b+ label allowed us to assign an inhibitory identity to cells, as GABA is the main inhibitory neurotransmitter in the zebrafish brain (Filippi, Mueller and Driever, 2014). Neurons displayed calcium dynamics similar to those previously reported (**Extended Data Fig. 1a** and (Bahl and Engert, 2020)). Unbiased clustering revealed three main functional cell types. Based on their dynamics, we named these cells motion integrator (MI), motion onset (MON), and slow motion integrator (SMI) neurons (**Fig. 1b, Extended Data Fig. 1a,b**, and **Methods**). Note that we previously used more descriptive labels based on the potential functional role of these cells within the evidence integration task (evidence integrator, dynamic threshold, and motor command neurons, respectively) (Bahl and Engert, 2020). Out of 15017 functionally imaged neurons, we found 618, 108, and 339 neurons of each type respectively, with the largest fraction located in the anterior hindbrain (62%: 360, 24, and 275 neurons of each type, respectively, **Extended Data Fig. 1c** and **Methods**) but also located in the pretectum (Pt), midbrain (Mb) and cerebellum (Cb) as well as other regions. Previous work suggests that the anterior hindbrain performs temporal integration of direction-selective whole-field motion inputs from the pretectum (Bahl and Engert, 2020). Compatible with these previous observations, we found that a large fraction of neurons in the anterior hindbrain were direction selective (DS) (75.6% of neurons had an absolute DS index>0.5, **Extended Data Fig. 1d)** with generally slower dynamics than their pretectum and midbrain counterparts (**Extended Data Fig. 1e**).

**Fig. 1.**
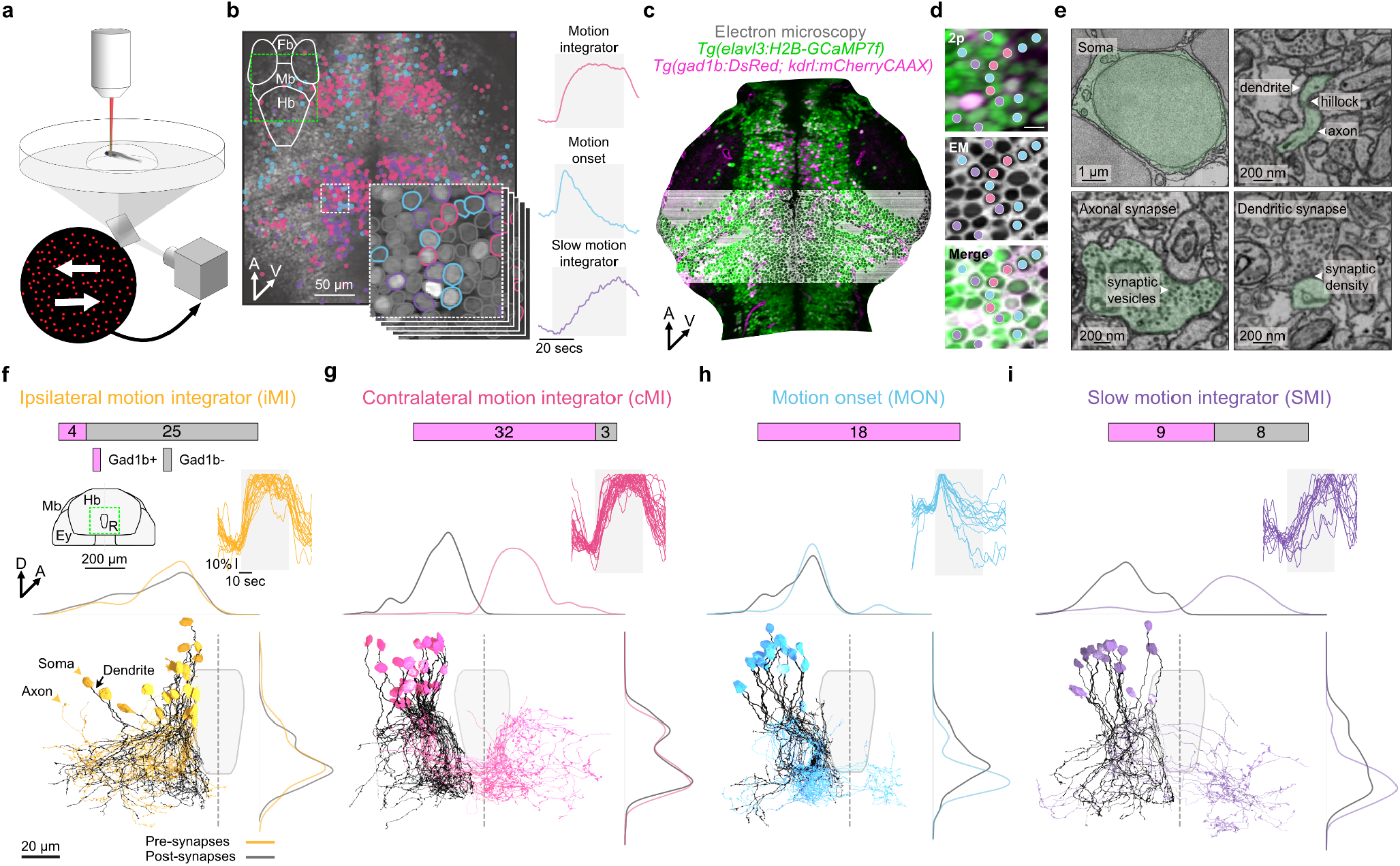
Correlative light and electron microscopy of neurons involved in motion evidence integration. **a**, Schematic of the two-photon imaging setup showing the bottom projection of coherently moving random dots. **b**, Left: spatial distribution of clustered neurons. The background grayscale image represents the average of the six functionally imaged planes. The top-left schematic shows coarse brain organization (dorsal-to-ventral view) and the location of the functionally imaged area (green dashed square). The snippets at the bottom right show individual imaging planes, with the top image showing an enlarged view of the region outlined by the white dashed square and also shown in **(d)**. Neurons are circled based on their functional types, while non-clustered neurons are circled in gray. Right: normalized K-means-based regressors used for neuron clustering. Stimulus consists of 0%, 100% (gray shaded area), and 0% coherence. **c**, Single plane of the two-photon image stack mapped to the EM dataset. Magenta shows inhibitory neurons (Gad1b+) and blood vessels (kdrl+). Green shows nuclear-localized GCaMP7f. Gray indicates the EM dataset. **d**, Detailed alignment between the two-photon imaging (2P) and EM datasets. Scale bar: 5 µm. Neurons from the clustered populations are indicated by colored dots. **e**, EM ultrastructural details of a reconstructed neuron. **f**, Structure-function relationship of functionally imaged and completely reconstructed neurons in coronal view. Top: distribution of neurotransmitter identity shown as Gad1b-positive (likely inhibitory, magenta) or Gad1b-negative (presumably excitatory, gray). Left: schematic showing coarse brain organization and location of the reconstructed neurons below (green dashed square) and outlines of the raphe (also shown below). Right: traces representing the normalized ΔF/F_0_ trial-averaged neuronal activity over time for motion in the preferred direction. Bottom: reconstructed ipsilateral motion integrator (iMI) neurons with somas and axons in orange and dendrites in black. Neurons reconstructed in the right hemisphere are mirrored to the left hemisphere for visualization purposes. The dashed line represents the midline. Surrounding plots are distributions of presynaptic (orange) and postsynaptic (black) synapses along the x (top) and y (right) axes. **g–i**, show identical representation for the other neuron types. All neurons and brain regions here and throughout the rest of the paper are registered to a reference brain atlas (Randlett *et al*., 2015). Abbreviations: Mb, midbrain; Hb, hindbrain; R, raphe; Ey, eye; A, anterior; D, dorsal; V, ventral. Neuron-type names, their abbreviations, and associated colors are used throughout the paper.

We next used the same larva, fixated it, and stained it for EM (Deerinck *et al*., 2010; Tapia *et al*., 2012; Hua, Laserstein and Helmstaedter, 2015) (**Methods**). To identify the coordinates of the block containing the relatively small and constrained region of the anterior hindbrain, we used sparse hindbrain electroporations (Boulanger-Weill *et al*., 2017) and c3pa-GFP photoactivation (Förster *et al*., 2018) in separate age-matched larvae (**Extended Data Fig. 2a,b**). Additionally, we performed X-ray computed tomography to ensure staining homogeneity and tissue integrity and to target the sectioning to specific neuronal circuits (**Extended Data Fig. 2c–g**).

We next collected 4010 coronal sections using an automated tape ultramicrotome (Hayworth *et al*., 2014) and performed multi-beam scanning EM (mSEM) at 4 x 4 x ~30 nm resolution (~416 x 265 x 112 µm, spanning from the medial midbrain to rhombomere 3, **Fig. 1c–e**), totaling 51.4TB of imaging data (**Methods**). Volumetric nanometer-precise alignment and connectome reconstruction were performed via a series of largely automated workflows, enabling high-quality neuronal morphology reconstruction ((Turner *et al*., 2022), **Methods**). To find single-cell correspondences between two-photon and EM imaging, we performed registration using manually placed corresponding landmarks ((Bogovic *et al*., 2016), **Methods**). This strategy yielded single-cell precision registration even in dense, low-contrast regions of the brain (Neuroglancer visualization, **Fig. 1d** and **Extended Data Fig. 2h**). Fluorescently labeled blood vessels, which were not used as landmarks for the registration, allowed us to independently confirm the quality of the mapping (**Extended Data Fig. 2h**). Finally, the EM and registered two-photon datasets were uploaded into a computational infrastructure enabling collaborative proofreading of morphology and connectivity of functionally identified neurons ((Dorkenwald *et al*., 2025), **Methods**).

### Structure-function relationship of functionally identified neurons

Within this dataset, we restricted our reconstructions to a defined region of the anterior hindbrain spanning rhombomeres 1–3 (corresponding primarily to Gad1b cluster 1-2 and Vglut2a cluster 2; ~100 × 100 × 100 µm), where approximately 40% of neurons display motion-integrating dynamics (Bahl and Engert, 2020). Within this volume, encompassing both hemispheres, we reconstructed 331 neurons, including 64 MI, 18 MON and 17 SMI and 63 myelinated neurons (including all reticulospinal neurons within the EM volume), as well as 56 input axons (Neuroglancer visualization, **Fig. 1f–i** and **Extended Data Fig. 3a–i**). To probe the connectivity of functionally identified neurons, we trained convolutional neural networks to automatically detect synaptic clefts and assign presynaptic and postsynaptic partner neurons ((Macrina *et al*., 2021), **Methods**).

All of these neurons were unipolar, with the axon emerging from a primary dendrite projecting ventrally, a structure reminiscent of invertebrate neurons (Dorkenwald *et al*., 2024; Schlegel *et al*., 2024). We observed distinct patterns among the different functional neuron types: most ipsilateral-projecting MI (iMI) neurons did not express Gad1b (Gad1b-, 25/29), while contralateral-projecting MI (cMI) neurons were mostly Gad1b positive (Gad1b+, 32/35). MON neurons were exclusively Gad1b+ and primarily contralateral-projecting (13/18), whereas SMI neurons exhibited a mix of neurotransmitter (9 Gad1b+ and 8 Gad1b-) and projection types: 5 ipsilateral-projecting Gad1b+ neurons and 12 contralateral-projecting neurons (**Fig. 1f–i**).

iMI neurons were generally positioned closer to the midline compared to other types, with overlapping axons and dendrites (**Fig. 1f** and **Extended Data Fig. 3e**). In contrast, cMI neurons were located more laterally, displaying clustered dendrites adjacent to the raphe, and had only a few axons leaving the volume (**Fig. 1g** and **Extended Data Fig. 3f**). MON neurons showed tightly clustered dendrites and axons (**Fig. 1h** and **Extended Data Fig. 3g**) with the majority of axonal synapses located on the ipsilateral side. SMI neurons had more dispersed dendrites and axons along both the dorso-ventral and rostro-caudal axes (**Fig. 1i** and **Extended Data Fig. 3h**) and were located more caudally. Interestingly, we did not find any statistically significant differences between presynaptic and postsynaptic synapse sizes or numbers across functionally identified cell types (**Extended Data Fig. 3j-k**).

### Distinct connectivity motifs across functionally identified neurons in the anterior hindbrain

To investigate connectivity motifs we first generated a connectivity matrix that included the 99 functionally identified neurons (**Fig. 2b)** together with myelinated neurons and axonal partners whose soma were located outside the EM volume. Distinct connectivity patterns emerged for each neuron type, both at the input and output levels (**Fig. 2b–j**). Importantly, connections described here were not indiscriminate. Indeed, reconstructed neurons arborize within the same neuropil, providing widespread opportunities for potential interactions, yet connected pairs exhibited significantly more similar response dynamics than non-connected visually responsive pairs occupying the same region (P < 10^-4^; Mann–Whitney U test). This relationship suggests that partner selection is functionally organized rather than random.

**Fig. 2.**
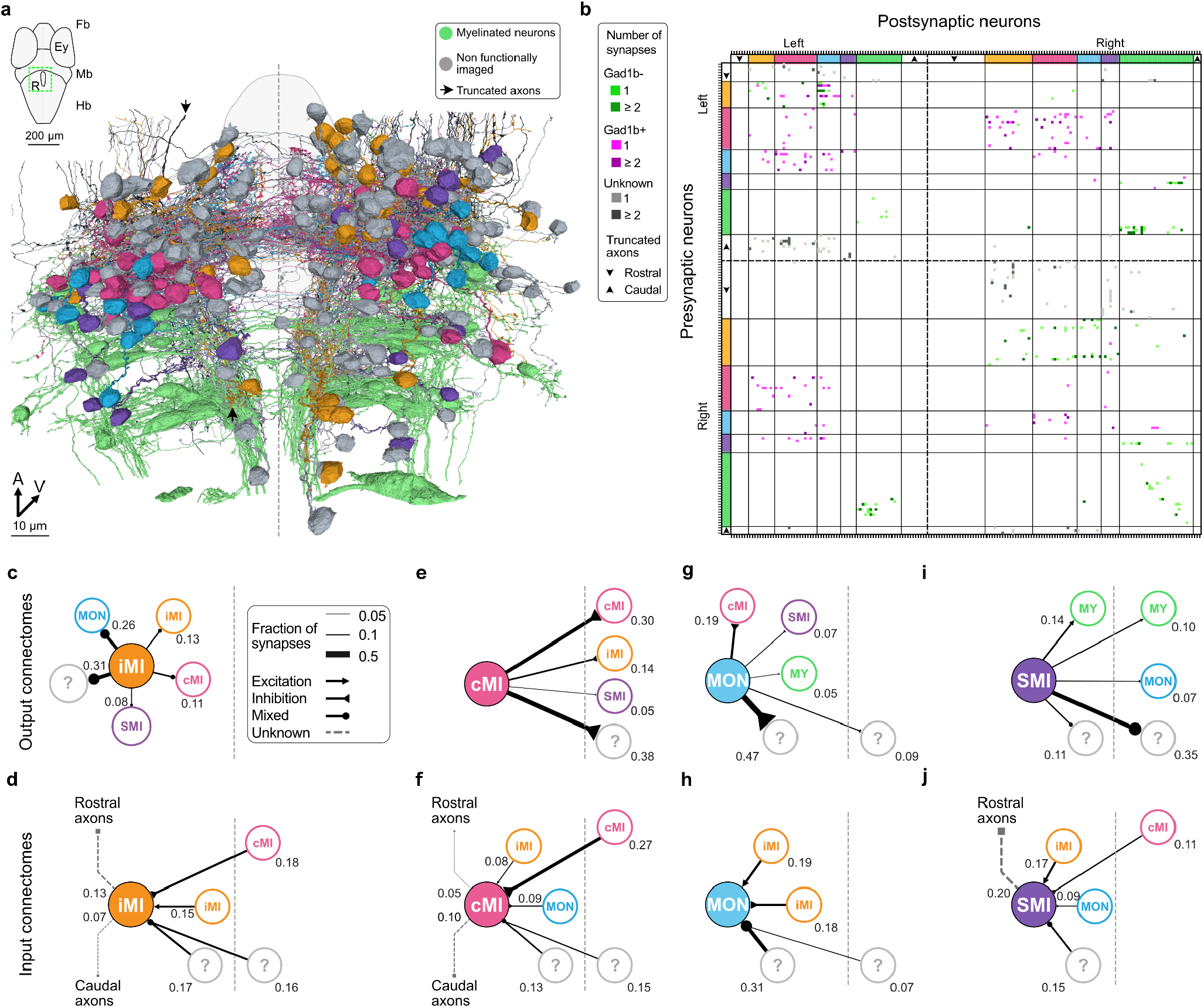
Connectomic analyses of functionally identified neuronal subtypes. **a**, Dorsal view of all neurons reconstructed in this study: 99 functionally imaged neurons, 63 myelinated neurons, 56 truncated axons and 169 non-functionally imaged neurons. Top left schematic shows coarse brain organization and location of the reconstructed neurons below (green dashed square) and outlines of the raphe (also shown below). **b**, Sparsely reconstructed connectivity matrix with neurons sorted according to their functional types and hemispheres (dashed black bars). Output (**c**,**e**,**g**,**i**) and input (**d**,**f**,**h**,**j**) cell-type connectomes extracted from the connectivity matrix. Connections accounting for less than 5% of the total synapse number are not shown. Question marks represent non-functionally imaged neurons. The dashed line indicates the midline. Abbreviations: Mb, midbrain; Hb, hindbrain; R, raphe; Ey, eye; A, anterior; V, ventral.

iMI neurons broadly targeted all functional types with a strong preference for MON, iMI and cMI neurons (26%, 13% and 11% of synapses, **Fig. 2c**). The substantial iMI-to-iMI component supports sparse recurrent connectivity within this neuronal subtype as previously hypothesized (Bahl and Engert, 2020). Additionally, a large fraction of the inputs originated from cMI neurons and from axons originating from the rostral end of the imaged EM volume (18 and 13% respectively, **Fig. 2d** and [Neuroglancer visualization](https://spelunker.cave-explorer.org/#!middleauth+https://global.daf-apis.com/nglstate/api/v1/6277423797108736) of an example connectome). cMI neurons primarily inhibited cMI and iMI neurons contralaterally (30% and 14% respectively, **Fig. 2e**) and received contralateral inhibitory inputs from cMI, ipsilateral inhibition from MON and ipsilateral excitation from iMI neurons (27%, 9% and 8% respectively, **Fig. 2f** and [Neuroglancer visualization](https://spelunker.cave-explorer.org/#!middleauth+https://global.daf-apis.com/nglstate/api/v1/6277423797108736) of an example connectome). These observations provide anatomical confirmation of predicted interhemispheric inhibition within the anterior hindbrain refining turning behavior by processing direction-selective inputs (Naumann *et al*., 2016).

MON neurons preferentially targeted cMI neurons (19%, **Fig. 2g**) and received mixed excitatory and inhibitory ipsilateral inputs originating from iMI (19 and 18%, respectively, **Fig. 2h** and [Neuroglancer visualization](https://spelunker.cave-explorer.org/#!middleauth+https://global.daf-apis.com/nglstate/api/v1/5891859952435200) of an example connectome). SMI neurons predominantly targeted caudally projecting neurons with myelinated axons (24%, both ipsilaterally and contralaterally, **Fig. 2i** and [Neuroglancer visualization](https://spelunker.cave-explorer.org/#!middleauth+https://global.daf-apis.com/nglstate/api/v1/5891859952435200) of an example connectome), confirming their role in motor execution, as previously proposed (Bahl and Engert, 2020). Consistent with this, one excitatory ipsilateral SMI neuron targeted three Rov3 reticulospinal neurons (see [Neuroglancer visualization](https://spelunker.cave-explorer.org/#!middleauth+https://global.daf-apis.com/nglstate/api/v1/5891859952435200) and **Extended Data Fig. 3i**), a subtype implicated in lateralized responses to whole-field motion (Orger *et al*., 2008). SMI neurons received excitatory input from iMI neurons (17%), contralateral inhibition from cMI neurons (11%), and ipsilateral inhibition from MON neurons (9%, **Fig. 2j**).

In summary, our sparse connectomic reconstructions in the hindbrain support a network model of bilateral inhibition across hemispheres mediated by cMI neurons, ipsilateral disinhibition driven by MON neurons, sparse recurrent connectivity within iMI neurons, and direct inputs from SMI neurons to reticulospinal neurons, which drive lateralized movements. To provide the broader scientific community with an online platform to browse neurons from connectomic analyses, we further developed an open-source atlas containing molecular labels and anatomic regions (Vohra *et al*., 2025) (see the [online atlas](https://fishexplorer.zib.de/lm/%7B%22stateId%22%3A%22671f916a0d5f0574aefcafac%22%7D)) to which we uploaded all reconstructed neurons and axonal segments (connected to functionally identified inputs and output partners).

### Two-photon-guided characterization of function, anatomy, and neurotransmitter identity

The initial two-photon imaging session was limited to a subset of imaging planes, resulting in a considerable fraction of reconstructed neurons lacking functional information. However, our reconstructions focused on a restricted anterior hindbrain region, where neuronal responses and morphologies can be organized into a relatively limited number of broad classes. This provided a favorable setting for investigating whether neuronal morphology could help predict the functional identities of neurons lacking direct recordings. We therefore sought to establish an independent reference dataset linking neuronal function, morphology, and neurotransmitter identity using light microscopy.

In a new set of experiments, we used two-photon excitation of photoactivatable GFP (Patterson and Lippincott-Schwartz, 2002; Förster *et al*., 2018) to reconstruct the anatomy of functionally characterized neurons (Kramer *et al*., 2019). We combined this approach with *in situ* RNA hybridization, HCR FISH (Choi *et al*., 2018), employing *gad1b* and *vglut2a* probes to selectively label GABAergic (inhibitory) and glutamatergic (excitatory) neurons, respectively (**Fig. 3a**). This strategy was applied across 47 larvae to generate a comprehensive multi-feature neuronal library that links functional dynamics, anatomy, and neurotransmitter identity—purely based on light microscopy experiments (**Fig. 3b–e**). Neuron functional types were determined through clustering analysis consistent with our FCLEM experiment (**Fig. 1b, Methods**). The resulting functional and morphological patterns closely resembled those observed in the FCLEM dataset (compare **Fig. 3b– e** and **Extended Data Fig. 4a–d** with **Fig. 1f–i** and **Extended Data Fig. 3e–h**), supporting the consistency of these observations across animals and imaging modalities.

**Fig. 3.**
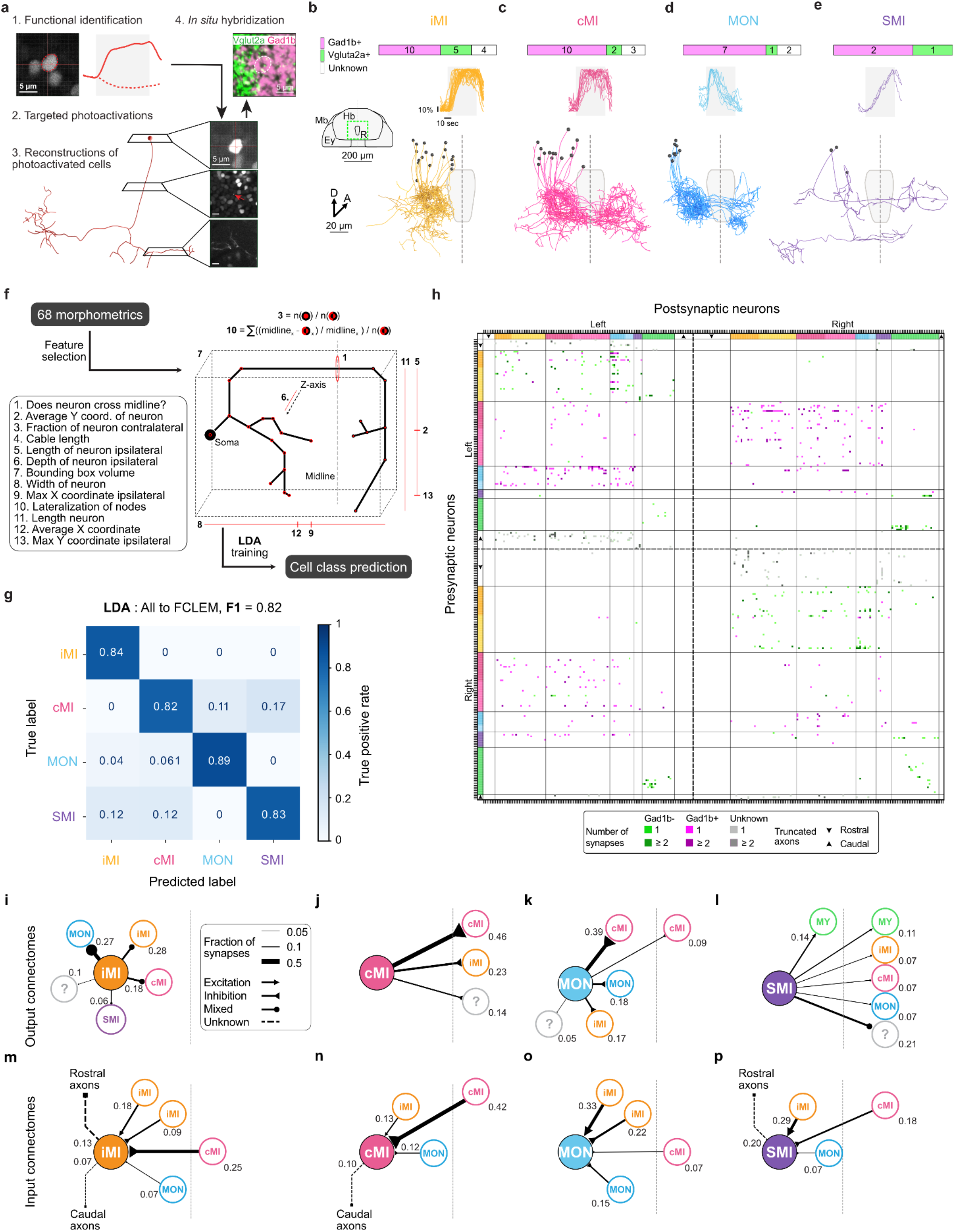
Single-cell photoactivation and functional type predictions across datasets. **a**, Schematic of the functional identification process followed by single-cell photoactivation, *in situ* hybridization, and manual morphological reconstruction. All scale bars correspond to 5 μm. **b**, Structure-function relationship of functionally imaged and photoactivated iMI neurons. Top: distribution of neurotransmitter identity shown as Gad1b-positive (magenta), Vglut2a-positive (green), or unknown (white). Left: schematic showing coarse brain organization (coronal view) and location of the reconstructed neurons below (green dashed square), as well as surrounding outlines of the raphe. Right: traces representing the normalized ΔF/F_0_ neuronal activity over time. Bottom: reconstructed iMI neurons with somas in black. The dashed line represents the midline. Abbreviations: Mb, midbrain; Hb, hindbrain; R, raphe; Ey, eye; A, anterior; D, dorsal; V, ventral. **c–e**, identical representation for other neuron types. **f**, Schematic of the feature selection used for the linear discriminant analysis (LDA) to predict functional cell types. **g**, LDA confusion matrix of true positive rate using FCLEM and photoactivated neurons as the ground truth and predicting FCLEM cells. The prediction performance F1-score for the entire matrix is indicated in the plot title. **h**, Sparsely reconstructed classifier-enhanced connectivity matrix of reconstructed EM neurons, sorted according to their functional types and hemispheres (dashed black bars). Neurons whose functional type has been predicted through LDA are indicated in lighter colors. Output (**i–l**) and input (**m–p**) cell-type connectomes extracted from the connectivity matrix. Connections accounting for less than 5% of the total synapse number are not shown. Question marks represent non-functionally imaged neurons that could not be classified by the LDA. The dashed line indicates the midline.

### Structure-to-function classification

Combining the photoactivation and FCLEM datasets yielded a reference library of 138 completely reconstructed neurons with known functional identities (43 iMI, 48 cMI, 27 MON, and 20 SMI). We used linear discriminant analysis (LDA) to identify combinations of morphological features that best distinguish the four functional classes. LDA is a statistical method used for dimensionality reduction and classification, identifying a linear combination of features that optimally separates two or more classes by maximizing variance between classes while minimizing variance within classes. Analysis of the model parameter contribution (**Methods**) suggested that only a few metrics had strong predictive power (**Fig. 3f–g** and **Extended Data Fig. 4e, Methods**). Applying a cross-validation holdout strategy (**Methods**), we achieved a classifier F1 score of 0.82 (**Fig. 3g**). This result indicates that neuronal morphology carries substantial information about broad functional identity within this restricted hindbrain region, although individual predictions remain subject to classification uncertainty.

We further tested the classifier’s performance with smaller datasets, using cells from either the photoactivation experiments or our FCLEM library. We still achieved predictive performance above 0.74 (F1-scores, **Extended Data Fig. 5a**). Remarkably, when training our classifier only on photoactivated cells and testing it on our FCLEM ground truth library, we maintained an F1-score of 0.76 (**Extended Data Fig. 5a**). We benchmarked our approach against other established methods including NBLAST (Costa *et al*., 2016; Li *et al*., 2017; Chen *et al*., 2020; Khosla *et al*., 2021; Choi, Kim and Hyeon, 2023; Dorkenwald *et al*., 2023) and found that our method consistently outperformed them in predictive accuracy (**Extended Data Fig. 5b–f**). These results demonstrate a partial correspondence between morphology and broad functional classes within the anterior hindbrain, while highlighting the need for additional validation before extending this approach to more heterogeneous neuronal populations in the zebrafish brain.

### Enhancing the FCLEM dataset

A direct application of this approach is the enhancement of our correlative light and EM dataset. Within this dataset, we identified 108 pre- and postsynaptic reconstructed unmyelinated neurons that we had not functionally imaged during our initial two-photon session (**Fig. 2a**). Using our trained classifier, we predicted the functional types of 78.7% of the non-functionally imaged neurons that were completely reconstructed (**Methods**), which considerably enriched our library with additional labels and allowed us to further populate the synaptic connectivity matrix of the anterior hindbrain (**Fig. 3h**).

To assess whether classification uncertainty affects the inferred population-level connectivity, we compared the fractions of output synapses from each presynaptic class onto each postsynaptic class between LDA-predicted and directly functionally identified neurons. These distributions were strongly correlated (Pearson’s *r* = 0.88; **Fig. 3i–p**). As an internal benchmark, we randomly divided the functionally identified neurons into two equally sized groups over 100 iterations and compared their output synapse fractions. These independent subsamples yielded a similar correlation (*r* = 0.83 ± 0.01 SEM). Thus, classifier-based augmentation preserves the major population-level connectivity motifs to a degree comparable to that observed between independently sampled subsets of functionally identified neurons.

### Morphological reconstructions and functional classification in a second whole-brain EM dataset

To test the generalizability of our classifier, we applied it to a WBEM volume (Petkova, 2025), which contains visual input regions absent from the FCLEM dataset (**Fig. 4a**,**b**). The WBEM dataset encompasses 800 × 450 × 250 μm from telencephalon to spinal cord and provides an ideal substrate for classifier-guided circuit reconstruction and long-range input reconstruction (**Fig. 4a–c**). We reconstructed 915 individual neurons connected by 1,643 manually annotated synapses, including neurons within the restricted anterior hindbrain region corresponding to our FCLEM analysis and long-range presynaptic partners ([Neuroglancer visualization](https://neuroglancer-demo.appspot.com/#!gs://fish1-release/assets/neuroglancer_states/202504/hindbrain_motion_integrator.json)). This reconstruction samples approximately 15–30% of neurons within the relevant anterior hindbrain clusters (**Extended Data Fig. 6a**), a reconstruction density which was shown to be sufficient to identify circuit motifs (Klinger *et al*., 2021). Before EM processing, the specimen, expressing Gad1b+ and VGluT2a+ labels, was imaged by confocal microscopy, enabling later assignment of neurotransmitter identity via single-cell co-registration with the WBEM dataset (**Fig. 4b**).

**Figure 4.**
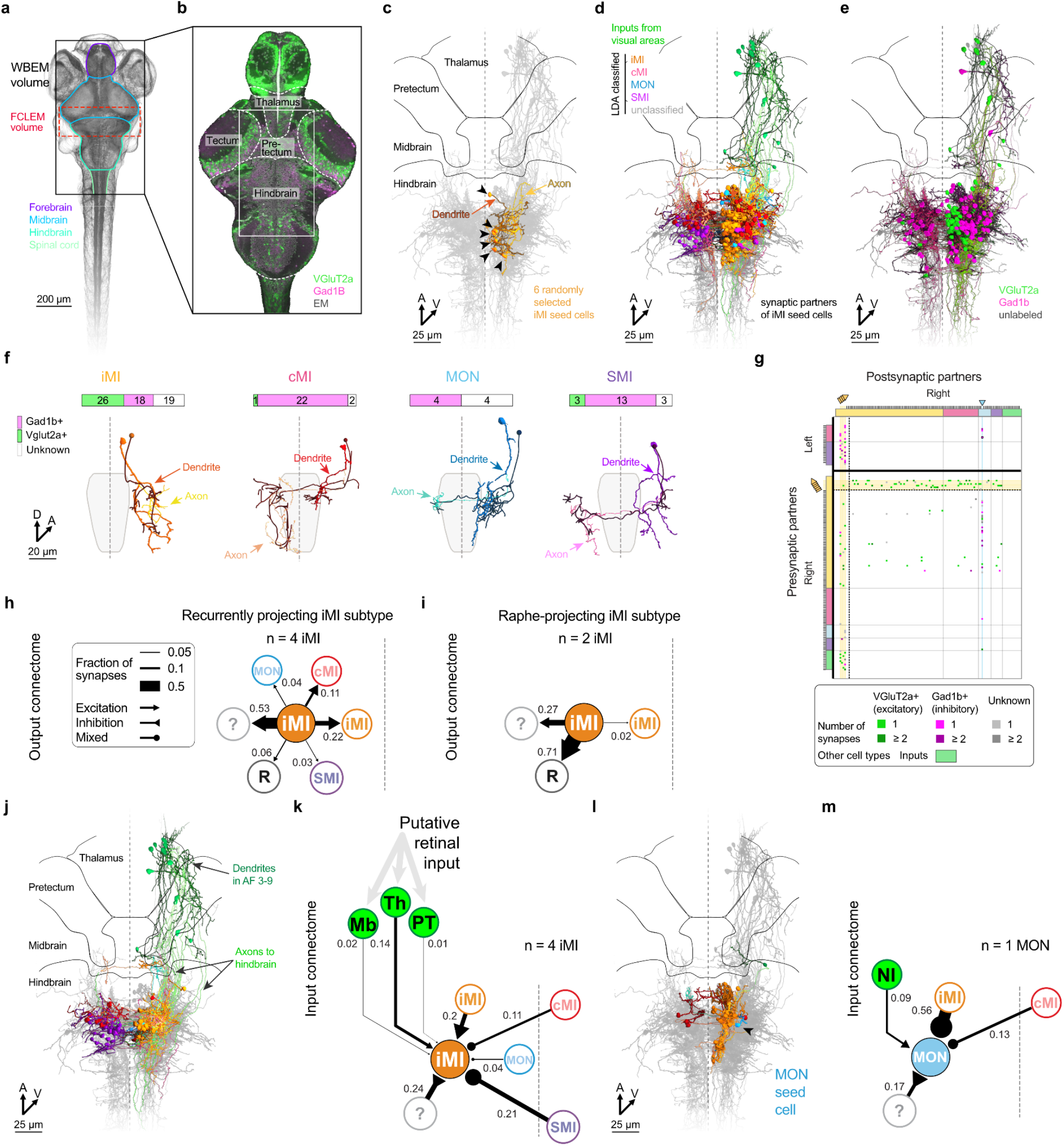
Connectivity of hindbrain integrator circuit reconstructed based on LDA classification in a second EM dataset containing the whole larval zebrafish brain. **a**, WBEM and FCLEM volumes. The other regions were not imaged. **b**, Single EM plane in gray with registered confocal imaging. Magenta shows inhibitory neurons (Gad1b+). Green shows excitatory neurons (VGluT2a+). **c**, Six randomly selected neurons classified as iMI using our classifier (**Fig. 3f**). Arrowheads indicate somas. **d**, Presynaptic and postsynaptic partners of four recurrently projecting iMI neurons, colored by predicted morphotype. **e**, Dorsal view of 215 reconstructed forebrain and anterior hindbrain neurons, colored by neurotransmitter identity. **f**, Top: neurotransmitter identity of all functionally classified WBEM neurons. Bottom: comparison between example photoactivated neurons from experiments in **Fig. 3a,b** and classified WBEM neurons. **g**, Connectivity matrix for LDA-classified WBEM neurons, organized by morphotype and hemisphere. Colored arrows and lines correspond to the densely reconstructed connectomes shown in **h–m. h–i**, Output connectivity diagrams of six iMI neurons, separated into four recurrently projecting and two raphe-projecting subtypes. All these neurons were considered excitatory. **j**, Presynaptic partners of four iMI neurons, showing long-range VGluT2a+ inputs from the ipsilateral thalamus, midbrain, and pretectum. **k**, iMI connectome extracted from the connectivity matrix. **l**, Presynaptic inputs to a MON neuron. **m**, MON connectome extracted from the connectivity matrix. Abbreviations: Mb, midbrain; Th, thalamus; PT: pretectum; NI: nucleus isthmi.

We first focused on six neurons classified as iMI as seed cells (four vGluT2a+, two unlabeled) (**Fig. 4c**). Their 273 outputs revealed 159 postsynaptic neurons, of which 115 could be assigned a functional type using our classifier. The resulting inhibitory/excitatory ratios and functional type distribution closely matched those observed in the FCLEM dataset (**Figs. 4d–f** and **1f–i**), demonstrating that the classifier generalizes across animals and EM datasets within this anterior hindbrain circuit. Analysis of iMI output connectomes revealed at least two distinct subpopulations: one displayed extensive iMI-to-iMI connectivity with frequent polysynaptic motifs (**Fig. 4g,h** and **Extended Data Fig. 6b)**; the other included two seed neurons that projected selectively to the dorsal raphe (**Fig. 4i**), suggesting a dedicated pathway for neuromodulatory feedback (Kawashima *et al*., 2016, 2024).

To investigate further the recurrent architecture of the network, we selected four iMI seed neurons from the first sub-population and exhaustively reconstructed their inputs and outputs. This comprises all first-degree presynaptic partners (N = 68), all first-degree postsynaptic partners (N = 113), and postsynaptic partners of first-degree output neurons (second-degree outputs; N = 825) (**Extended Data Fig. 6c**). The second-degree outputs revealed extensive recurrence: 67 neurons were rediscovered in previously reconstructed network layers, corresponding to 36% of neurons in the combined input, seed, and first-degree output populations. Further examination of the connectivity among the seed neurons, their first-degree inputs, and first-degree outputs revealed a densely interconnected central integrator cluster (**Extended Data Fig. 6d**). This cluster exhibited connection reciprocity and clustering coefficients comparable to those reported for densely reconstructed connectomes in *Drosophila, C. elegans*, the zebrafish oculomotor integrator, and mouse primary visual cortex (Lin, 2024). We also observed similar rediscovery rates in the FCLEM volume (40.54% and 22.83% for left and right iMI populations, respectively), indicating that substantial multi-step connectivity is present in both specimens despite differences in reconstruction coverage.

Using the classifier to interpret the reconstructed network, we found that the first-degree presynaptic partners (40/68 were assigned a functional type via our classifier) originated from two principal sources. First, local anterior hindbrain inputs, with connectivity patterns consistent with the recurrent excitation and bilateral inhibition motifs observed in the FCLEM dataset (**Figs. 4j** and **3m**). Second, long-range excitatory ipsilateral inputs from multiple rostral brain regions. Surprisingly, most of the inputs arose from the thalamus with fewer contributions from the midbrain and pretectum (**Fig. 4j,k**). Many input neurons extended dendrites into retinal arborization fields, consistent with direct input from retinal ganglion cells (Kubo *et al*., 2014; Naumann *et al*., 2016; Svara *et al*., 2022). Although the pretectum was previously considered the primary source of motion-selective input (Naumann *et al*., 2016), recent evidence indicates that the thalamus can also convey visual information (Zhao *et al*., 2026). Consistent with this broader view, our findings identify multiple visual areas that innervate iMI neurons.

Although we do not directly measure this specificity in our FCLEM and WBEM datasets, our complementary results suggest the presence of wide-field direction-selective neurons in the thalamus and other input regions. To probe this area for such activity, we performed two-photon calcium imaging targeting this and adjacent regions during random dot motion stimulation (**Extended Data Fig. 7a–b**). As expected, direction-selective responses were observed in the tegmentum, optic tectum, and pretectum, but also in the thalamus. Moreover, the majority of WBEM input cells were located within 20 μm of a direction-selective neuron when mapped to a common reference brain (**Extended Data Fig. 7b**). We also quantified the rise time constants for these input regions (**Extended Data Fig. 7c-f**) and found that they exhibited faster dynamics than the anterior hindbrain, consistent with its role in temporal integration. Taken together, these results suggest that iMI neurons receive long-range, direction-selective inputs from multiple visual processing regions, with the thalamus representing a potentially important source.

Having established the long-range visual inputs to iMI neurons, we next asked whether MON cells might also receive direct synaptic input from early visual brain regions. We identified a MON-classified neuron in the anterior hindbrain and reconstructed all its presynaptic partners (**Fig. 4l,m**). We found a strikingly similar local input structure to that of functionally identified MON cells in the FCLEM dataset (**Fig. 3o**). Taken together, these results confirm that MON neurons are targeted by local iMI neurons relaying whole-field visual flow from rostral visual areas.

In summary, the independent WBEM dataset revealed local connectivity motifs consistent with those observed in FCLEM, while more extensive tracing uncovered higher-order recurrent connectivity and long-range inputs from rostral visual regions. These findings support the reproducibility of the identified circuit organization across specimens and illustrate the value of morphology-based functional classification within this restricted anterior hindbrain region, where functional and morphological diversity is relatively limited.

### A connectome-constrained recurrent neuronal network model reproduces and predicts cell type dynamics

We next sought to investigate whether the experimentally reconstructed circuit provides a sufficient anatomical substrate compatible with the different temporal dynamics observed in the anterior hindbrain. To this end, we designed a bilateral recurrent neural network model directly derived from the measured neuron-to-neuron connectome and the measured excitatory and inhibitory labels (**Fig. 3h**). The network receives leftward and rightward coherence-dependent visual motion inputs, targeting the corresponding ipsilateral hemispheres (**Fig. 5a**). We restricted excitatory external sensory input to neurons receiving anatomically reconstructed rostral projections (green squares in the input mask, **Fig. 5b**). The experimentally identified connections then defined the binarized masked connectivity arrangement within the network. Although EM reconstructions reveal which neurons are synaptically connected, they do not directly allow inference of their effective physiological synaptic strengths. We therefore employed synaptic weight optimization to fit model dynamics to the experimentally measured ones for each functional cell type.

**Figure 5.**
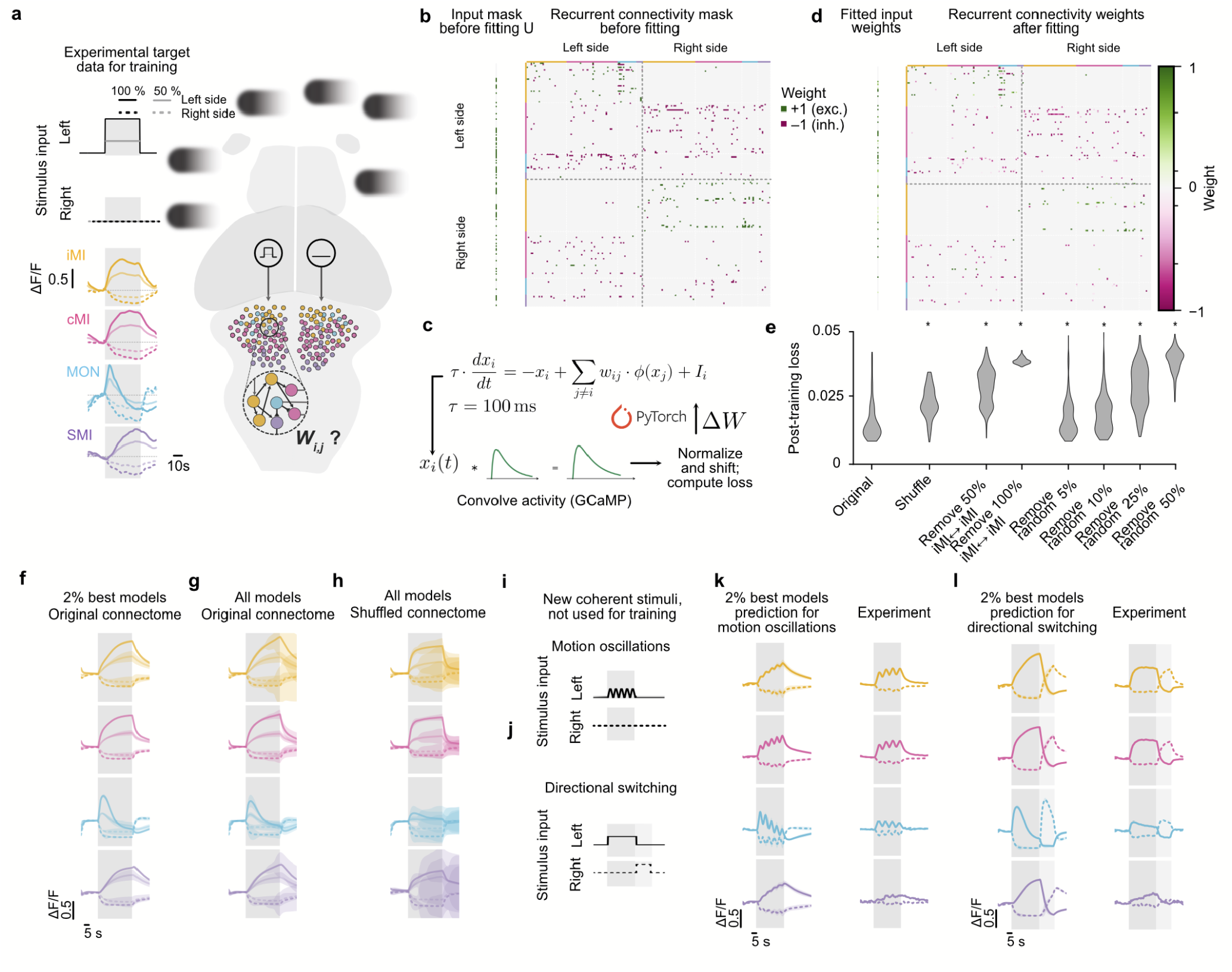
A connectome-constrained neural network model of the anterior hindbrain. **a**, Experimentally measured target functional cell-type activity used for fitting. Gray-shaded regions indicate periods of motion. Responses to both coherence levels are derived from the traces in **Fig. 1f–i** by scaling and adding different noise patterns (**Methods**). For leftward whole-field motion (illustrated by leftward drifting coherent dots), only the left side of the model hindbrain receives stimulus input signals. For rightward motion, data is mirrored (not shown). We seek to estimate the weight matrix in the anterior hindbrain that can generate such dynamics as well as the input weight vector. **b**, Binarized recurrent connectivity matrix with neurotransmitter labels directly taken from experiments (**Fig. 3h**). The input weight vector (left green squares) was masked based on whether input cells with rostral projections were present or not. **c**, Fitting pipeline. All neurons have a time constant of 100 ms. Activity is convolved with a calcium-response kernel following published kernels (**Methods**), allowing for error (loss) comparisons with experimental imaging data. **d**, Example fitted input weight vector and recurrent connectivity weight matrix. Weights are displayed in a logarithmic color scale. **e**, Loss function distributions (300 model runs each) for the original experimentally constrained recurrent connectivity matrix without any modifications, with shuffled recurrent connectivity mask, and with different degrees of recurrent connection removals (**Extended Data Fig. 8d,e** for example masks). Note that the removal of 100% iMI to iMI connections corresponds to around 5 % removal of random connections across the recurrent connectivity mask. Statistical comparisons are based on two-sided Mann-Whitney U test (each p < 0.001), comparing loss at the end of training in networks with and without structural manipulations (Cohen’s d is 0.26 and 0.55 for models with random 5% and 10% connections removed respectively, >1.2 in all other cases). **f**, Dynamics of the best 2% models with original constraints (mean across models ± STD; N = 6 models). **g**,**h**, Same as in (f), but for all N = 300 model fitting runs, comparing the original network with shuffled controls. The median correlation coefficient between model readout and population targets is r_m_=0.91 with interquartile range r_IQR_=0.02 using original connectome constraints, and r_m_=0.74 with r_IQR_=0.12 using shuffled connectivity masks. **i**,**j**, New, previously untested, motion stimuli used to challenge the fitted models: Leftward-biased motion coherence that oscillates between 0 % and 100 % at 0.16 Hz (i) and continuous leftward drift (100 %) that abruptly switches direction (100 % in the opposite direction, j). The lighter gray shaded area indicates the period after the switch. **k**,**l**, Same as in (**a**,**f**) but for the new coherent motion stimuli shown in (i,j), respectively. Left columns are model predictions, and right columns indicate experiments. Same models as in (f). N = 1 fish in (a) and N = 6 fish in (k,l). Correlation coefficients between model readout and population targets were in the range [0.79, 0.84] for motion oscillations, and in [0.89, 0.93] for directional switching.

All neurons in the model follow a standard exponential rate model with a rate time constant of 100 ms, which is within a biologically plausible range (Grama and Engert, 2012; Vishwanathan *et al*., 2024). In training and testing phases, simulated unit activity was first convolved with a GCaMP response kernel and then compared with population-averaged calcium responses across the four functional classes, hemispheres, and stimulus coherence levels (**Fig. 5c, Methods**). The result was a network with optimized weights for the input vector and the recurrent connectivity matrix that preserved the exact experimental connectivity and neurotransmitter distribution (**Fig. 5d**). We repeated the optimization 300 times, allowing us to compute distributions of loss values (**Fig. 5e**). We then validated the dynamics and model-to-model variability for the best 2% model outcomes (**Fig. 5f**). These networks’ output activity was highly correlated with the experimental traces shown in Fig. 5a (median Pearson’s r_m_=0.90, with interquartile range r_IQR_=0.02, across models). Despite the fast intrinsic neuronal time constant of 100 ms, the optimized networks reproduced the distinct temporal dynamics of the four functional classes, including the slow multiple-second-long accumulation and decay characteristic of motion integrating neurons. These slow dynamics are not due to the GCaMP response kernel convolution as MON cells displayed their characteristic brief onset dynamics. Although the model was fitted to population average responses, dynamics were also present in individual model neurons for each functional cell type and across different model weight matrix outcomes (**Extended Data Fig. 8a**,**b**). This result was confirmed by computing the distribution of correlation coefficients between single-cell activity traces and their experimental counterpart (**Extended Data Fig. 8c**). We next assessed whether the neurons with the highest response amplitude exhibited dynamics consistent with the experimental recordings. To this end, we recomputed the distribution of correlation coefficients using only neurons whose activity amplitude fell within the highest quartile of their respective populations. These neurons likewise exhibited target-like dynamics (**Extended Data Fig. 8c, Methods**).

To determine whether our results depend on the experimentally observed connectivity organization, we compared networks trained using the recurrent connectivity mask from the original connectome with networks trained using shuffled recurrent connectivity masks. We designed the shuffling step such that it preserved the overall connectivity rate in the network and all neurotransmitter labels (**Methods, Extended Data Fig. 8d**). Shuffling the recurrent connectivity mask significantly reduced fit quality (**Fig. 5e**, higher loss function values). To compare dynamics between conditions, we plotted the average results from all fitted models (**Fig. 5g,h**), showing that shuffled connectomes led to worse matching dynamics with larger variability across model outcomes (**Fig. 5g,h**).

We next investigated the contribution of specific connectivity motifs. Our anatomical reconstruction revealed recurrent excitatory connections among ipsilaterally projecting iMI neurons, suggesting a potential role of this motif in the temporal integration of sensory input during prolonged visual motion stimulation. To test this hypothesis, we selectively removed either 50% or 100% of iMI to iMI connections from the anatomical connectivity mask before optimization (**Fig. 5e** and **Extended Data Fig. 8e**, for 100% removal). Both manipulations impaired the abilities of the fitted networks to reproduce the experimentally observed dynamics, as measured by significant increases in the loss distributions. Notably, removing a comparable fraction of randomly selected connections across the whole network (5% overall corresponds approximately to 100% within iMI) produced substantially smaller impairments (**Fig. 5e** and **Extended Data Fig. 8g)**, indicating that the contribution of recurrence within the iMI population to fitting quality cannot be explained simply by a reduction in overall connectivity across the network. Removing larger fractions of connections progressively reduced model performance, showing that also the general connectivity within the network plays an important role (**Fig. 5e**). Together, these results identify the connectivity among iMI neurons as an important circuit motif for supporting the dynamical features within the reconstructed network.

Next, to determine whether our modeling benefited from the LDA-based connectome enhancement, we repeated the optimization without assigning predicted functional identities to neurons that had not been functionally imaged. All reconstructed neurons and their anatomically observed connections were retained, but neurons lacking direct functional recordings remained functionally unconstrained, contributing to network dynamics without being assigned response targets or contributing directly to the loss function. Importantly, despite the absence of these functional constraints, the network retained its ability to reproduce slow integration dynamics. Even though such an arrangement reduces the number of functional constraints on the network and may thus simplify the problem, the resulting loss function distribution increased significantly (**Extended Data Fig. 8h**). Plotting the dynamics of the unlabeled cells after fitting – within their predicted LDA-based category – showed that they overall matched the predicted cell type dynamics (**Extended Data Fig. 8i**), corroborating that connectivity and functional features are correlated within the anterior hindbrain. We also tested a network that only contained the original non-enhanced FCLEM dataset where cell numbers are smaller (compare **Figs. 3h** and **2b**). As expected, optimization results showed considerable reductions in fitting quality (**Extended Data Fig. 8h**). Together these tests demonstrate the benefit of our classification approach to enhance connectomes with incomplete functional labels.

To assess whether the optimized models could generalize beyond the stimuli used during training, we designed two novel random-dot-motion stimulation paradigms (**Fig. 5i,j**). The first stimulus consisted of oscillating motion coherence, probing how the different neuronal populations filter temporally varying sensory cues (**Fig. 5i**). The second one introduced abrupt changes in motion direction, allowing us to better probe the dynamics of bilateral interactions across the network (**Fig. 5j**). We selected the top 2% of models based on their performance on the training data and evaluated their predictions for the test stimuli. The preselected models predicted distinct responses across the four functional populations. During oscillating motion coherence, iMI and cMI neurons exhibited slow integration dynamics with superimposed oscillations, MON neurons displayed their prominent transient onset feature with subsequent oscillating decay, and SMI neurons increased their activity more gradually (**Fig. 5k**). During directional switching, models predicted a relatively rapid redistribution of activity between hemispheres, accompanied by transient responses in MON neurons (**Fig. 5l**). We subsequently tested these predictions experimentally using functional calcium imaging and found that our models captured the principal response dynamics across neuronal populations for both stimulus types (**Fig. 5k,l**). Thus, our constrained model networks can generalize across new stimuli that have not been included during optimization.

In summary, our connectome-constrained model fitting demonstrates that the experimentally reconstructed connectivity arrangement provides an anatomical substrate compatible with the dynamics observed in the anterior hindbrain, including slow integration of visual motion over several seconds. Connectivity perturbations in the model identify a specific computational contribution of iMI-to-iMI recurrent connections, linking an experimentally observed connectivity motif to temporal integration. The fitted model further predicts responses to novel visual stimuli, showing that our EM-based circuit model can generate new experimentally testable hypotheses.

## Discussion

Linking neuronal dynamics to synaptic organization remains a major challenge in vertebrate neuroscience. Here, we combined functional imaging with electron microscopic reconstruction to identify the neuronal morphologies and connectivity patterns associated with processing noisy visual motion in the larval zebrafish anterior hindbrain. We identified four broad functional and morphological classes interconnected through recurrent excitation, interhemispheric inhibition, and ipsilateral inhibitory pathways (**Fig. 1-3**). Together, these findings provide direct anatomical evidence for several circuit motifs previously proposed from functional imaging (Naumann *et al*., 2016; Bahl and Engert, 2020) and establish a framework for investigating their computational contributions.

Recurrent excitation among motion-integrating neurons was previously proposed as a mechanism supporting slow temporal integration in the anterior hindbrain (Bahl and Engert, 2020). Our reconstructions provide direct anatomical evidence for recurrent excitation, with approximately one quarter of iMI output synapses targeting other iMI neurons across two independent EM datasets (28% in FCLEM and 22% in WBEM, **Fig. 3-4**). Denser reconstruction in the WBEM dataset further revealed substantial local recurrent connectivity, with 36% of neurons rediscovered across three successive network layers (**Fig. 4**). To investigate the computational relevance of this architecture, we developed a network model constrained by the experimentally measured neuron-to-neuron connectivity (**Fig. 5**). Although EM reconstruction reveals anatomical connections, it does not directly determine their effective physiological synaptic strengths. We therefore used machine learning techniques to optimize connection weights within the observed anatomical architecture to investigate whether it provides a substrate compatible with the measured temporal dynamics. Despite fast intrinsic neuronal time constants, the resulting networks reproduced slow population dynamics and predicted responses to novel visual stimuli. Importantly, selectively removing iMI-to-iMI connections substantially degraded the recovery of slow integration dynamics, whereas removing an equivalent number of connections anywhere in the network had a much smaller effect. These findings identify recurrent excitation as a particularly influential component of temporal integration within the anatomically constrained model, although the physiological synaptic weights underlying integration *in vivo* remain to be determined.

Beyond recurrent excitation, our reconstructions revealed additional connectivity motifs associated with bilateral interactions and sensorimotor processing. In particular, inhibitory projections from cMI neurons onto contralateral iMI populations provide anatomical support for bilateral inhibitory interactions previously proposed from functional imaging (Naumann et al., 2016; Bahl and Engert, 2020). This organization is also consistent with opponent inhibitory motifs implicated in selective signal amplification and decision-making in mice (Zagha, Ge and McCormick, 2015; Kuan *et al*., 2024). We additionally identified distinct input and output pathways involving MON and SMI populations, as well as long-range projections from several rostral visual regions, including the thalamus. These observations extend previous circuit models by revealing additional pathways through which visual information can reach motion-integrating populations and be transmitted toward downstream motor circuits.

An important simplification of our analysis is the grouping of neurons into four broad classes, which does not capture the full diversity of neuronal response properties and connectivity within each population. Indeed, our reconstructions already reveal substantial anatomical heterogeneity among iMI neurons, including locally recurrent populations and neurons projecting selectively to the dorsal raphe (**Fig. 4**). Similar diversity exists within cMI and SMI populations, potentially reflecting finer functional subdivisions that remain unresolved by our current classification. Recent connectome-based analyses of the zebrafish oculomotor integrator have demonstrated that neurons participating in the same broad computation exhibit considerable diversity in their sensitivity to eye position and temporal response dynamics (Vishwanathan et al., 2024). Likewise, neuronal variability within the motion-processing populations identified here could reflect differences in integration timescales, sensitivity to motion coherence, or other stimulus properties.

Our modeling framework further simplifies neuronal diversity by optimizing network responses against population-averaged calcium dynamics. Although individual anatomical connections are retained, the model does not explicitly address how differences in single-neuron response properties or trial-to-trial variability contribute to sensory integration. Such variability may be particularly relevant in the anterior hindbrain, where neurons exhibit alternating left–right activity during spontaneous exploration (Dunn *et al*., 2016) and integrate visual inputs in a state-dependent manner to bias reorienting swim bouts during phototaxis (Wolf *et al*., 2017). These findings indicate that visual information processing depends not only on sensory inputs but also on the ongoing state of the network. Future studies combining behavioral recordings, denser anatomical reconstructions, voltage imaging, electrophysiological recordings, transcriptomic analyses, and targeted perturbations will be essential to determine how cellular and synaptic properties shape these dynamics. Recent studies in *Drosophila* illustrate how additional physiological constraints can be incorporated into connectome-based models to investigate circuit computations (Ammer *et al*., 2023; Lappalainen *et al*., 2024; Shiu *et al*., 2024).

Finally, the connectomes and analysis tools presented here are openly available through an integrated zebrafish brain atlas (Vohra *et al*., 2025), complementing existing vertebrate connectomic resources (Svara *et al*., 2022; Petkova, 2025). Although our reconstructions sample only a fraction of the underlying circuit, they provide a foundation for progressively linking neuronal function, synaptic connectivity, and computation. More broadly, our results demonstrate how combining functional measurements with anatomically constrained modeling can identify connectivity features relevant to population dynamics while providing a framework for investigating the neuronal diversity underlying vertebrate sensorimotor computations.

## Methods

### Zebrafish

For the correlative light and EM experiment (FCLEM dataset, **Figs. 1** and **2**), we generated a triple transgenic *Tg(elavl3:H2B-GCaMP7f*^*+/+*^, *gad1b:dsRed*^*+/-*^ (Satou *et al*., 2013), *kdrl:mCherryCAAX*^*+/-*^ (Fujita *et al*., 2011)) larva obtained by crossing adult transgenic *Tg(elavl3:H2B-GCaMP7f, gad1b:DsRed)* and *Tg(elavl3:H2B-GCaMP7f, kdrl:mCherryCAAX)* lines in nacre background, *mitfa*^*−/−*^ (Lister *et al*., 1999), yielding high GCaMP7 fluorescence. The *Tg(elavl3:H2B-GCaMP7f)*^*u344Tg*^ transgenic line was generated using standard methods (Dowell *et al*., 2024). We raised small groups of 20–30 larvae in filtered fish facility water in Petri dishes (9 cm in diameter) on a 14 h light, 10 h dark cycle at a constant 28 °C. From 4 dpf onwards, we fed larvae with paramecia once per day. The experiment was performed on 7-dpf larvae.

For sparse cell electroporations (**Extended Data Fig. 2a**), we incrossed *Tg(elavl3:H2B-GCaMP7f)*^*u344Tg*^ lines and selected offspring based on green fluorescence. For the area photoactivation (**Extended Data Fig. 2b**), we crossed *Tg(alpha-tub:c3pa-GFP*^*+/-*^)^*a7437Tg*^ (Bianco *et al*., 2012) with *Tg(elavl3:H2B-GCaMP6s*^*+/-*^)^*jf5Tg*^ (Freeman *et al*., 2014) adults and selected offspring for green fluorescence. Animals were 7 dpf and 6 dpf old, respectively.

For the combined functional recordings, single-cell photoactivation, and HCR FISH staining experiments (**Fig. 3**), we outcrossed *Tg(alpha-tub:c3pa-GFP*^*+/-*^; *elavl3:h2b-GCaMP6s*^*+/-*^) and *Tg(alpha-tub:c3pa-GFP*^*+/-*^)^*a7437Tg*^ adults.

For the imaging experiments in **Figs. 4** and **5**, we used incrossed *Tg(elavl3:GCaMP8s)*^*kn6Tg*^, a newly generated line expressing *GCaMP8s* in neurons. To generate the line, we injected the Tol2 vector transgene construct (*Tol2-elavl3:GCaMP8s* (Zhang *et al*., 2023)), obtained from Janelia Research Campus, and transposase RNA into 1–4-cell-stage embryos. We then isolated transgenic lines by screening for high expression of bright green fluorescence in the central nervous system in the next generation.

For the experiments shown in **Figs. 3**–**5**, larvae were raised in E3 water on a 14h light, 10 h dark cycle at a constant 28 °C without feeding. Animals were 5 dpf old.

All experiments were approved by the Harvard University and Konstanz University committees on the use of animals in research and training.

### Visual stimuli

All visual stimuli during functional imaging consisted of ~1,000 dots (2 mm in diameter) moving at 1.8 cm s^−1^, projected (60 Hz, P300 Pico Projector, AAXA) from below through mildly light-scattering parchment paper. Dots were shown in red on a black background to minimize interference of the visual stimulus with the functional GCaMP imaging. To ensure that the animals did not track individual dots to detect motion, each dot had a short lifetime (200 ms mean) and stochastically disappeared and immediately reappeared at a random location within the arena. One stimulation trial consisted of 40 s of 100% coherent motion either drifting leftward or rightward, interleaved by 20 s of 0% coherence before and after. For the FCLEM experiment (**Fig. 1**), we also presented moving sine gratings drifting leftward or rightward, but did not include these stimulus trials in our analysis. For the imaging experiments in **Fig. 5i–l**, we presented animals with 30 s of 100 % coherence, with motion either running continuously or abruptly switching direction after 15 s. Stimuli across trials were interleaved with 40 s of 0% coherence. For oscillating-coherence trials, motion coherence varied between 0% and 100% at 0.16 Hz in a fixed direction. In all experiments, stimuli were shown in a random sequence.

### Two-photon calcium imaging and analysis

For the FCLEM experiment, the larva was fully embedded in a drop of agarose (UltraPure Low Melting Point Agarose, Invitrogen), at ~35 °C in the center of a Petri dish (9 cm in diameter, VWR). After the agarose had solidified, the dish was filled with fish facility water and transferred into the measurement chamber of a custom-built two-photon microscope, operated by custom-written Python 3.7-based software (PyZebra2P). We used a femtosecond-pulsed laser (MaiTai Ti:Sapphire, Spectra-Physics) equipped with a set of x/y-galvanometers (Cambridge Technology), a 20x infrared-optimized objective (XLUMPLFLN, Olympus) to scan over the brain, and two photomultipliers (green and red) amplified by two current preamplifiers (SR570, Stanford) to collect fluorophore emissions. We used frame acquisition rates of around 1 Hz. Laser power was measured to be ~13 mW at the specimen. Such power levels did not interfere with behavior in previous tail-free imaging experiments (Bahl and Engert, 2020). During functional imaging, the laser was tuned to 950 nm to collect GCaMP7f fluorescence. We sequentially imaged six planes spaced at 12 μm, centered on the anterior hindbrain region (336 x 336 μm at 0.48 x 0.48 μm resolution). In each plane, we showed from 2 to 9 visual stimulation trials. Immediately after the experiment, we anesthetized the fish by replacing the water with a 0.015% MS-222 (Sigma-Aldrich) solution in fish facility water. This procedure enabled us to acquire an additional motion artifact-free high-resolution stack at 1020 nm excitation (420 x 420 x 90 μm at 0.6 x 0.6 x 2 μm resolution) with simultaneous collection of GCaMP7f and DsRed fluorescence. The total imaging session lasted ~5 hours.

Activity recordings of each plane were first motion-corrected using NoRMCorre (Pnevmatikakis and Giovannucci, 2017) and temporally averaged. We used a convolutional neural network segmentation approach (StarDist) to perform high-accuracy segmentation of the nuclei in each plane (Weigert and Schmidt, 2022). Calcium dynamics for each nucleus mask were then extracted as ΔF/F_0_ traces using Python numpy array slicing.

K-means clustering of functional cell types was performed on pooled activity traces from the FCLEM experiment (**Fig. 1**) and all reconstructed neurons from our targeted photoactivation experiments (**Fig. 3**). The elbow method (Umargono, Suseno and S. K., 2019) suggested 4 functionally distinct clusters. These clusters were further isolated by two successive K-means clustering steps with 3 and 2 target clusters as parameters. The first iteration isolated MON dynamics, fast motion integrator dynamics (MI-1), and a third, slower group. The second iteration performed on this slower population revealed an additional motion integrator group (MI-2) as well as a slow motion integrator group (SMI). Thus, our clustering approach yielded 4 neuronal clusters with distinct temporal dynamics. For all cells in each cluster, we computed the average rise time until neuronal activity reaches 90% of the maximal response, revealing a clear separation of timescales across clusters (**Extended Data Fig. 1b**, only showing FCLEM cells). We then computed 3 final regressors by pooling neurons belonging to the MI-1 and MI-2 clusters and averaging the activity of all neurons belonging to each clustered population. Neurons in the FCLEM recording were then assigned to one of the 3 functional types by computing the correlation coefficient with individual regressors. Neurons whose correlation coefficient with a regressor was larger than 0.8 and that had a response of at least 20 % ΔF/F_0_ were assigned to the corresponding functional type. All other neurons remained unassigned. The resulting clusters were spatially distinct and interpretable: for example, SMI neurons were located more caudally and, as demonstrated in previous work (Bahl and Engert, 2020), were preferentially recruited at the onset of motor tail bouts, in line with their proposed role as output-stage units of the integrator circuit.

Direction selectivity indexes (**Extended Data Fig. 1d**) were computed using standard methods (Boulanger-Weill *et al*., 2017):

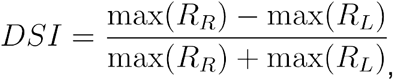

where *max(R*_*R*_) and *max(R*_*L*_) are the maximal ΔF/F_0_ response values during the entire rightward and leftward stimulation. Negative responses were set to 0.

For the imaging experiments presented in **Extended Data Fig. 7**, and **Fig. 5i-l**, we used our newly generated fish line *Tg(elavl3:GCaMP8s)*^*kn6Tg*^. Raw imaging data was processed as previously described and neurons were segmented using cellpose3 (Stringer and Pachitariu, 2025) on time-averaged imaging stacks, using the following cyto3 model parameters: cell diameter = none; follow threshold = 0.7; cellprob threshold = –0.4; lower = 0; upper = 90.0, which generated very good segmentation results across the brain (**Extended Data Fig. 7a**). We computed ΔF/F_0_ time traces from the resulting masks. In the imaging experiments with the new oscillatory and switching stimuli presented in **Fig. 5i-l**, we also displayed 100 % coherence continuously moving rightward or leftward (not shown). This configuration allowed us to use the same data processing, regressors, and cell type classification strategy as for our other imaging experiments (**Figs. 1** and **3**). We then averaged all labeled cells per group and fish to obtain response dynamics.

In the imaging experiments presented in **Extended Data Fig. 7**, individual neuron centroids were registered to the reference brain using ANTs (Avants, Tustison and Johnson, 2015; Randlett *et al*., 2015). Next, we identified responsive cells by performing a one-sided t-test between the peak responses during stimulus and during pre-stimulus baseline lasting 10s for each cell. If either the left-motion or right-motion test had a p-value smaller than 0.005, we considered the cell responsive. We computed DSI as described previously and classified a cell as DS if its absolute DSI > 0.33. We then computed, for each DS neuron, the time until neuronal activity reached 90% of the maximal response during motion onset. We split the data by brain region in **Extended Data Fig. 7c**. We extracted these brain regions from the mapzebrain atlas ((Kunst *et al*., 2019); downloaded on 19-06-2025 from https://mapzebrain.org) and registered them to the reference brain using ANTs. We compared whether the distribution of rise-times of each region (dorsal thalamus, pretectum, optic tectum, and tegmentum) was lower than the hindbrain distribution using a one-sided Mann-Whitney U test with Holm-Bonferroni correction.

### Sparse electroporations and area photoactivations

Sparse electroporations were performed as in (Boulanger-Weill *et al*., 2017). Briefly, 4-dpf larvae were embedded in a droplet of agarose and anesthetized with 0.015% MS-222 (Sigma-Aldrich) in fish water. The agarose next to the anterior hindbrain was removed using a microsurgical blade. The electroporations were performed under a fixed-stage upright microscope equipped with a long working distance objective (10x UPlanFL N 10X/0.30na, Olympus). The silver wire microelectrode and bath electrode were mounted on micro-manipulators, with the microelectrode holder containing an additional side port to apply air pressure with a syringe. Borosilicate glass capillaries (1.2 mm outer diameter, 0.68 mm inner diameter, with filament by World Precision Instruments) were pulled using a pipette puller (PC-10, Narishige, set at 68/58 °C) to obtain a 1 μm tip diameter. Pipettes were filled with 2 μL of *cmv:lyn-tdTomato* plasmid DNA solution and inserted into the anterior hindbrain of larvae. The lyn tag enables high-efficiency targeting of the red fluorescent protein to the cell’s membrane and is thus ideal for studying neuronal morphology (Kunst *et al*., 2019). Following optimized protocols (Zou, Friedrich and Bianco, 2016), two or three stimulation bursts (around 0.5 s per burst, consisting of 2 ms pulses at 200 Hz at 28 V) were delivered using a square pulse stimulator (Sd9, Grass Technologies) and monitored using a digital oscilloscope (TDS 2014, Tektronix). After the electroporations, larvae were freed from the agarose and kept isolated for 2 days. At 6 dpf, we then performed two-photon functional imaging of GCaMP6s (see previous section) to measure the response dynamics of the labeled cells to random dot motion. For subsequent morphological imaging of electroporated cells, we then anesthetized the fish and imaged at 1030 nm to collect dual-color z-stacks at high spatial resolution (0.46 x 0.46 x 1.5 μm) of both GCaMP and td-Tomato. Morphological imaging data were filtered (3D Gaussian blur, sigma 1.0 in x, y, and z) in Fiji (Schindelin *et al*., 2012). Labeled neurons were then semi-automatically traced with the open-source reconstruction software neuTube (Feng, Zhao and Kim, 2015), using default settings and optimal node resampling. We then saved cells as .swc files. Using the GCaMP6s channel, we then generated a transform per larva, allowing us to map reconstructed electroporated cells to a reference brain (Randlett *et al*., 2015).

For area photoactivation (**Extended Data Fig. 2b**), fish were anesthetized with 0.015% MS-222 (Sigma-Aldrich), and c3pa-GFP was photoactivated using cycles of 40 short pulses of 200 ms at 1 Hz at a laser wavelength of 750 nm, following established protocols (Kramer *et al*., 2019). After 1–3 initial trial pulses to check whether any adjacent cells were photoactivated and to readjust the region of interest if necessary, the whole protocol consisting of 15 cycles was executed, with five-minute intervals between two activation cycles. Photoactivation was targeted over a 50 x 40 μm region containing neurons with motion integration dynamics in the anterior hindbrain.

### Sample preparation for the EM imaging

For the FCLEM experiment, we used a protocol modified from (Deerinck *et al*., 2010; Tapia *et al*., 2012) to enhance extracellular space preservation, which improves synapse detection (Pallotto *et al*., 2015) and permits single-cell resolution imaging using X-ray tomography. Unless noted, all steps were performed at room temperature (RT). Immediately after the two-photon stack acquisition, the larva was still anesthetized and embedded in agarose. We replaced the water with a dissection solution (64 mM NaCl, 2.9 mM KCl, 10 mM HEPES, 10 mM glucose, 164 mM sucrose, 1.2 mM MgCl_2_, 2.1 mM CaCl_2_, pH 7.5) supplemented with 0.02% tricaine (Hildebrand *et al*., 2017). We then cut small slits in the agarose to expose the eyes and performed bilateral enucleations to enhance extracellular space preservation and heavy metal staining. We used a custom-made hook that was carefully inserted behind the eyes to prevent brain damage. The larva was immediately transferred to a fixation solution at 4°C (2.5% glutaraldehyde, 0.1M cacodylate buffer supplemented with 4.0% mannitol at pH 7.4). Cacodylate buffer: 0.3M sodium cacodylate, 6 mM CaCl2, pH 7.4. To improve the fixation, the tissue was rapidly microwaved (cat. no. 36700, Ted Pella, equipped with a power controller, steady-temperature water recirculator, and cold spot) in the fixative solution (this step lasted <5 min after initial transfer into fixative).

The microwaving sequence was performed as in (Tapia *et al*., 2012): at power level 1 (100 W) for 1 min on, 1 min off, 1 min on then to power level 3 (300 W) and fixed for 20 s on, 20 s off, 20 s on, three times in a row. Fixation was then continued overnight at 4°C in the same solution. The following day, the sample was then washed again in 0.5x cacodylate buffer (3 exchanges, 30 min each) before osmication (2% OsO_4_ in 0.5x cacodylate buffer, 90 min). After a quick wash (< 1 min) in 0.5x cacodylate buffer the sample was reduced in 2.5% potassium ferrocyanide in 0.5x cacodylate buffer for 90 min then washed with double-distilled H_2_O (3 exchanges, 30 min each) and then incubated with 1% (w/v) thiocarbohydrazide (TCH) in double-distilled H_2_O (and filtered with a 0.22 μm syringe filter before use) for 45 min to enhance staining (Hua et al., 2015). Due to poor dissolution of TCH in water, the solution was heated at 60 °C for ~90 min with occasional shaking before filtering and then placed at RT for 5 min before the incubation step. The sample was then washed with double-distilled H_2_O (3 exchanges, 30 min each) before the second osmication (2% OsO_4_ in double-distilled H_2_O, 90 min) and then washed again (3 exchanges, 30 min each). Then, en-bloc staining was performed overnight using 1% uranyl acetate in double-distilled water at 4°C. The solution was sonicated for 90 min and filtered with a 0.22 μm syringe filter before use. Steps involving uranyl acetate were performed in the dark. The following day, samples were then washed with double-distilled H2O (3 exchanges, 30 min each).

Next, the samples were dehydrated in serial dilutions of ethanol (25%, 50%, 75%, 90%, 100%, 100% for 10 min each step) and then in propylene oxide (PO) (100%, 100%, 30 min, each step). Infiltration was performed using LX112 epoxy resin with BDMA (21212, Ladd) in serial PO dilution steps, each lasting 4h (25% resin/75% PO, 50% resin/50% PO, 75% resin/25% PO, 100% resin, 100% resin). Samples were mounted in fresh resin in a mouse brain support tissue (Hildebrand *et al*., 2017) with the head exposed to facilitate cutting and prevent the sample from sinking to the bottom of the resin molds. Mouse tissue was fixed using standard procedures (Fang *et al*., 2018) and then cut into 2–3 mm wide cubes, which were pierced using a puncher (0.75 mm, 57395, EMS) to insert the larva. The cubes were stained along with fish samples using the protocol described above, except that the uranyl acetate overnight step was performed at RT. The samples were then cured with support tissue for 3 days at 60 °C. For all steps, a rotator was used. Aqueous solutions were prepared with water passed through a purification system (Arium 611VF, Sartorius Stedim Biotech). The protocol lasted 5 consecutive days, including surgery, fixation, staining, and resin embedding, followed by 3 days of resin curing.

### X-ray computed tomography

To achieve high contrast X-ray and mSEM imaging, we sequentially optimized OsO_4_ staining duration, X-ray imaging, and voltage, and performed deep learning-based X-ray reconstruction (**Extended Data Fig. 2e,f**). The OsO_4_ staining was optimized first in previous batches of samples to yield the mSEM staining protocol described previously. The imaging voltage and post-processing were optimized in the main sample described in this study. In total, this sample was imaged for ~300 hours, which did not interfere with mSEM imaging. For each sample, the block was trimmed to expose the head of the fish for imaging (Xradia 520 Versa, ZEISS). To assess the imaging quality, we computed a modified version of Michelson’s contrast that was robust to noise at multiple sections along the main axis of the larvae:

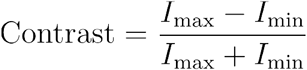

Where I_max_ and I_min_ are the average intensity values over the 80^th^ and below the 20^th^ percentile, respectively. We also computed the signal-to-noise ratio (SNR) at the same locations according to (Joy, 2002). X-ray tomography stacks were post-processed using a deep neural network product included in the Xradia software.

### Sample sectioning, post-staining, and mSEM image acquisition

After X-ray imaging, the block was re-embedded in resin and cured for 3 days at 60 °C and trimmed into a diamond shape (Hildebrand *et al*., 2017). 4010 sections ranging from 30 to 35 nm were automatically collected on carbon-coated tape using a custom tape collection device (ATUM) (Hayworth *et al*., 2014) mounted to a commercial ultramicrotome. The tape was cut into strips and deposited on 30 silicon wafers, which were post-stained to enhance membrane contrast using a protocol adapted from (Shapson-Coe *et al*., 2024). Briefly, each wafer was first plasma-treated for 30 seconds (operating pressure of 6 x 10^-1^ mbar, plasma current of 10 mA) to increase its hydrophilicity and then immediately stained with 4% uranyl acetate for three minutes, rinsed with double-distilled H_2_O for three minutes three times, air-dried, stained with 3% lead citrate for three minutes, rinsed and air-dried as before and stored overnight under vacuum.

Wafers were then mounted on the 61-beam mSEM stage (MultiSEM 505, ZEISS), and each section’s position was determined using a reflected light microscope to guide the high-resolution imaging. Imaging was performed at 4 x 4 nm pixel resolution using secondary electron emission with a dwell time of 400 ns per pixel. The quality of each section was assessed using previously described methods (Shapson-Coe *et al*., 2024).

### mSEM image alignment, stitching, rendering, and neuronal morphology proofreading

EM image alignment, cell segmentation, and synapse detection/assignment were outsourced to Zetta AI. Convolutional neural networks were trained to detect defects and classify locations that contain tissue for each montaged section. Self-supervised convolutional neural networks were used to generate dense displacement fields between sections (Popovych *et al*., 2024). The elastic energy in the set of all displacement fields was then hierarchically minimized to produce an aligned image. An “image mask” was created to mark defects (folds, wrinkles, missing sections) and locations that were classified as misaligned, preventing these artifacts from degrading downstream segmentation and synapse detection. Additional convolutional neural networks were trained to segment cells, detect synaptic clefts, compute their sizes, and assign presynaptic and postsynaptic objects from the cell segmentation at each cleft (Macrina *et al*., 2021). Synapses were identified based on the combined presence of (i) synaptic vesicles, (ii) darker synaptic clefts, and (iii) presynaptic density. Autapses arising from segmentation errors were automatically removed before constructing the connectivity matrices. After this filtering, synapses detection had a true positive rate of 89.5% and an inter-user agreement of 95% (two expert users, 1011 synapses corresponding to **Fig. 3h**). Cell segmentation, synapse detection, and assignment used the aligned image with locations in the “image mask” set to black. Finally, cell segmentation was ingested into the ChunkedGraph proofreading system so that morphological errors could be manually corrected, and synapses were ingested into an annotation table for consistent analysis (Dorkenwald *et al*., 2025).

For the neuronal reconstructions used in **Fig. 1** and LDA training, two expert tracers independently proofread the same cells during rounds 1 and 2, followed by a third fine-tuning round performed by one tracer, following a strategy similar to (Dorkenwald *et al*., 2022). Four representative neurons, one from each functional class, illustrate this procedure (**Extended Data Fig. 3a–d**). Synapses were retrieved from each reconstruction state; relative to the final agreement state, mean synapse-retrieval F1 score was 0.81 after round 1 and 0.91 after round 2, with mean tracing times of 28.8 and 46.8 min per neuron, respectively. Other neurons and axons underwent one or two proofreading rounds.

### Multimodal, reference brain registrations and cell type determination

Matching of neuronal identities from the 2P functionally imaged planes to the EM volume was performed in two successive manual steps using the Fiji plugin Bigwarp (Bogovic *et al*., 2016). We first manually registered functional imaging planes to a higher z-resolution two-photon image stack (z-steps = 2 μm, containing green and red fluorescence) using 844 manually selected landmarks. The SNR of this stack was enhanced using content-aware image restoration (Weigert *et al*., 2018). We then registered this higher resolution two-photon image stack to a downsampled mSEM stack (0.512, 0.512, 0.480 μm/pixel) using 771 manually selected landmarks. This EM stack was downloaded from Neuroglancer using Cloudvolume (Silversmith *et al*., 2022). These two registration steps generated two deformation fields, which were successively applied to the 2P functionally imaged planes using Bigwarp (in linear deformation mode) and manually checked exhaustively. The segmented neuronal masks were also deformed using these two steps (in nearest neighbor mode). Deformed planes of green and red fluorescence (8-bit image stacks) and segmented masks were uploaded to Neuroglancer using CloudVolume, enabling collaborative reconstruction and cell type determination of the neurons of interest. Given the large number of cells in the EM dataset (approximately 62745), it was challenging to comprehensively quantify the error rate across the entire sample. Instead, to ensure reliability of cross-modality mapping (two-photon to EM), we visually inspected every neuron included in this study, carefully evaluating the quality of local alignment. When misalignments were detected, we applied locally refined transformations to correct residual errors before inclusion in downstream analyses.

To determine whether neurons were inhibitory, we manually traced a mask surrounding the neurons’ nuclei in the green channel and computed the average corresponding red fluorescence. Neurons with values greater than 10 % of the maximum red signal (25 on an 8-bit scale) were considered inhibitory. Regions containing k-means clustered neurons (**Extended Data Fig. 1c**) were determined by automatically registering the 2P functionally imaged planes to the z-brain reference brain (Randlett *et al*., 2015) using ANTs (Avants, Tustison and Johnson, 2015). For regional annotations, we downloaded anatomical masks from the mapZebrain atlas platform (Kunst *et al*., 2019) and mapped them into the z-brain coordinate system (Randlett *et al*., 2015) using ANTs (Avants, Tustison and Johnson, 2015).

Finally, neurons traced in Neuroglancer were downloaded using Navis (Schlegel *et al*., 2025) and automatically registered to the z-brain reference brain (Randlett *et al*., 2015). To register cells from our downsampled mSEM stack to the z-brain reference brain, we manually selected 207 landmarks in the mSEM stack and the *Tg(elavl3:H2B-RFP)* confocal stack in the z-brain coordinate system to generate a bridge transform via Fiji Bigwarp. All neurons presented in this paper have been registered to the z-brain reference brain (Randlett *et al*., 2015) and are available on our online platform.

### Two-photon photoactivation and neurotransmitter identity of functionally identified cells

We selected larvae for green fluorescent nuclear expression and embedded animals in a droplet of low-melting-point agarose in a Petri dish with a diameter of 6 cm. The fluorescence of the H2B-GCaMP6s makes the identification of c3pa-GFP difficult – at this stage, c3pa-GFP is not yet photoactivated and thus very dim. To enable screening for c3pa-GFP, we initially performed a brief activation experiment in the two-photon microscope, labeling a small region in the dorsal rostral tectum. We only continued with the animals that showed some sign of tectal photoactivation patterns after a maximum of 5 rounds of photoactivation. Morphologies originating from this area largely remained local and did not overlap with the photoactivated neurons in the hindbrain. In a few cases, neurons descended into the anterior hindbrain neuropil, coming close to the photoactivated cells, but did not interfere with our tracing. To acclimate larvae to the experimental conditions and minimize motion artifacts and plane shifts during imaging, we initially presented animals for 15 minutes with random dot motion before starting the functional characterizations.

To quickly identify cell types, we then imaged a single plane in the anterior hindbrain for 15 – 30 minutes while presenting random dot motion. Using a custom-written Python script, we aligned recordings to the onset of motion and averaged stacks across trials. Using Fiji (Schindelin et al., 2012), we defined a small movable region of interest with the size of a cell to manually explore the stack for neuronal dynamics matching the known cell type dynamics (Bahl and Engert, 2020). This analysis step took less than 5 minutes per fish.

For each identified cell, we then conducted targeted photoactivation at 760 nm. Using a custom-written scanning waveform generator in PyZebra2P, we used a small activation spiral (less than 0.5 µm in diameter), focused onto the center of the nucleus of the cell. Due to the point spread function of our microscope (~1 µm along the horizontal plane, and ~4 µm along the vertical axis), we were able to activate c3pa-GFP in the cytosol without affecting neighboring neurons. Following recent protocols (Kramer *et al*., 2019), we used short activation pulses of 200 ms in length and 100 ms interpulse intervals. The required laser power at the specimen and the pulse counts for effective photoactivation varied widely across fish. Some fish only required as few as 20 pulses (ca. 6 seconds) at 5 mW to activate cells, whereas other fish often required up to 1000 pulses (ca. 5 min) at 8 mW. We attribute this difference to variations in expression levels, as some experimental fish are heterozygous and some fish are homozygous for c3pa-GFP.

Following successful photoactivation, we anesthetized the fish by replacing the water in the Petri dish with a 0.015% MS-222 solution. After 15 minutes of anesthesia, we then imaged a high-resolution stack (100 planes, 0.4 µm x 0.4 µm x 2 µm; 90 s per plane) of the hindbrain and midbrain to capture as much of the photoactivated cell’s neurite morphology as possible. This step took around 2.5 hours per animal.

To determine the neurotransmitter identity of the labeled cell, we performed HCR RNA-FISH (Choi *et al*., 2018) on the same fish immediately after the imaging session. To this end, we cut out a small block of agarose containing the anesthetized fish and directly fixed the sample in ice-cold 4% PFA for 12–16 hours overnight at 4 °C. We then followed recent protocols optimized for zebrafish larvae (Shainer *et al*., 2023). We used probe sets against *gad1b* (probe set size 30, #NM_194419.1) and *vglut2a* (*slc17a6b*, probe set size 20 #NM_001128821), with B1 and B2 amplifiers, respectively. We used the B1-546 amplifier fluorophore and the B2-405 amplifier fluorophore, thus fluorescently labeling *gad1b*-positive cells in red and *vglut2a*-positive cells in blue.

After the procedure, we placed the still agarose-embedded fish into the same two-photon microscope where it had been imaged before. The PFA fixation and HCR RNA-FISH protocol considerably reduced the fluorescence of a photoactivated c3pa-GFP cell. Cells were still identifiable because they were slightly more fluorescent than neighboring non-photoactivated neurons. Neurites in the photoactivated cells were, however, not visible anymore after the procedure. To further validate the identity of the cell, we used landmarks such as the arrangement of neighboring nuclei or neuropil structures with distinguishable distances to the labeled cells. We acquired two separate high-resolution volume scans around the photoactivated cell body. Our custom-built microscope can only simultaneously image green and red fluorescence or green and blue fluorescence, with a manual replacement of a filter cube in the collection optics path between imaging sessions. As we acquired green fluorescence in both sessions, we could identify our target cells in both imaging stacks to extract the expression levels of both *gad1b* and *vglut2a* within the cytosolic ring around the nucleus. Labels were largely mutually exclusive, allowing us to confidently assign neurotransmitter identity based on which label was stronger. We labeled cells where both *gad1b* and *vglut2a* signals were indistinguishable from background fluorescence as ‘undefined’.

The custom-designed two-photon microscope used for these experiments was largely similar to the one explained above, used for the FCLEM experiment. However, this system used a tunable MaiTai® DeepSee (Spectra-Physics) laser source with a built-in pre-chirp unit, a Nikon 25x Objective (N25X-APO-MP), and a more sensitive Hamamatsu GaAsP PMT (H16201P-40-S3) for the green channel. These features significantly improved the imaging quality of green signals and later facilitated the reconstruction of photoactivated neurons. Reconstructions were performed in Fiji using the SNT semi-automated tracing tool (Arshadi *et al*., 2021).

### Zebrafish-optimized NBLAST probability matrix

We used NBLAST (Costa *et al*., 2016) to compute a matrix of morphological similarity of cells within our FCLEM dataset (**Extended Data Fig. 5f**). One of the major functions of the NBLAST algorithm is the 2-dimensional scoring matrix (10 x 21), which provides a raw score of whether two neurons belong to a similar cell type. The rows of the scoring matrix represent the dot product of the tangent vectors, whereas the columns represent the range of distances in micrometers. The default matrix is derived from a dataset of *Drosophila* neurons and is sensitive to several factors, including dataset type, neuron size, preprocessing methods, and the volume analyzed. To match the algorithm to our data, we computed a zebrafish-optimized scoring matrix using the FCLEM dataset. The training of such a matrix requires matching and nonmatching sets of neurons (Costa *et al*., 2016). We used our four broad functionally and morphologically distinct cell types (iMI, cMI, MON, SMI) as matching sets. Non-matching sets of neurons are drawn randomly from our FCLEM dataset. The consistency of the matrix is assured by using the n-fold cross-validation (n=4). The detailed method and further advancements are provided in an upcoming atlas paper (Vohra *et al*., 2025).

### Morphology-based prediction of neuronal functional types

With the unique availability of two ground truth datasets of structure and function relationship (**Fig. 1** and **Fig. 3**), we sought to employ and train a classifier to predict a functional cell type based on anatomical structure. Such a classifier will be useful for predicting functional response properties of cells in anatomical libraries for which functional imaging has not been performed, including existing EM reconstruction databases. To this end, we first mapped all reconstructed neurons to a common reference atlas (Randlett *et al*., 2015), using custom-written Python scripts and ANTs (Avants, Tustison and Johnson, 2015). For neurons originating from the FCLEM experiment (**Fig. 1**), we applied the same Bigwarp-based generated ANTs transforms that we used to map cell bodies into the reference brain (see section above). For neurons originating from the two-photon guided photoactivation experiments (**Fig. 3**), we used the h2b-GCaMP6s channel to generate a transform between each stack and the corresponding reference stack.

Reconstructed neurons from the FCLEM experiment were downloaded as high-resolution geometry (.obj) files. These files are simple text files containing a list of surface vectors describing volumetric meshes, allowing us to apply our ANTs transforms to map geometries into the z-brain coordinate system. Using the *skeletor* package and the TEASAR algorithm (Sato *et al*., 2000) in Python, we skeletonized .obj files to 1.5 µm spacing and saved results as .swc files. This procedure ensured that all neurons originating from different experiments and modalities ended up in the same coordinate system with the same spatial tracing resolution, a prerequisite for the development of our classifier.

As our NBLAST matrix does not show a clear separability of cell types, we decided to build our classifier using a feature-based approach. To this end, we exhaustively extracted 68 morphometrics from each skeletonized neuron (**Extended Data Table 1**). All features were normalized by subtracting the mean and scaling to unit variance across cells. To identify the most relevant of these features for predicting functional cell types, we first selected feature subsets of different sizes. To this end, we conducted feature ranking with recursive feature elimination using *sklearn*.*feature_selection*.*RFE* in Python with “AdaBoostClassifier” as the learning estimator. For a given target feature subset size (1 – 68), the algorithm starts at the maximum number of features (68). It then uses the specified estimator to evaluate the importance of features by fitting the model and then iteratively removing the least significant feature until it reaches the desired target feature subset size. For each resulting feature subset, we then extracted a predictive F1-score of how these features could be used to classify cell types. Specifically, we split our cells (including data from both the FCLEM and photoactivation experiments) in a ratio of 70% train and 30% test. As a classifier algorithm, we chose Linear Discriminant Analysis (LDA) with solver ‘lsqr’ (least squares solution). Performing the LDA training and prediction step 100 times with different randomly selected training and test sets allowed us to compute the predictive F1-score as a function of the feature subset size (**Extended Data Fig. 4e)**. We then selected the feature combination with the highest predictive performance as the final feature set (**Fig. 3f**). We used this feature subset for all further cell type classifications.

Many cells identified through our pre- and post-synaptic connectomic reconstructions had their soma in regions that were not covered by our 2P multi-plane imaging session. To compute the performance of our classifier when predicting the functional type in such cases, we performed a ‘leave-one-out-cross-validation’ step on our ground truth datasets. To assess the quality of the classifier using maximal and cross-modality training data, we combined both the FCLEM and photoactivation experiments. We iteratively chose one of our known FCLEM cells and trained our classifier on all remaining ground truth cells. Comparing the classifier prediction of the left-out FCLEM cell with its true functional type allowed us to compute a confusion matrix, indicating the probability that a functional prediction of a given FCLEM cell is correct (**Fig. 3g**). We finally repeated the training on all available ground truth cells (data from the FCLEM and photoactivation experiments) to generate the final classifier used to predict the functional type of FCLEM and WBEM cells that were not functionally imaged (**Figs. 3h, 4c–d**). Our trained cell type classifier should also be applicable to enhance other morphological resources, containing anatomical reconstructions of neurons in the larval zebrafish anterior hindbrain.

Notably, the LDA classifier will place any target cell, even when it looks completely different from the training dataset, into one of our defined 4 categories. To rule out such incorrect predictions, we identified outliers within the predicted cells’ morphometric features using an isolation forest algorithm (*sklearn*.*IsolationForest*) and the local outlier factor (*sklearn*.*LocalOutlierFactor*). We trained these outlier removal methods with the morphometric features of our functionally identified cells from both the FCLEM and photoactivation experiments.

We compared our morphology-based neuronal type prediction method to previously established approaches (Li *et al*., 2017). To do so, we computed persistence vectors and their sampled versions (Li *et al*., 2017) for each neuron using *navis*.*persistence_vectors* and trained a linear discriminant analysis classifier (*sklearn*.*LinearDiscriminantAnalysis*) to predict functional types (**Extended Data Fig. 5b,c**). Similarly, we calculated form factors (Choi, Kim and Hyeon, 2023) for each neuron using *navis*.*form_factor* and trained another linear discriminant analysis classifier to predict functional types (**Extended Data Fig. 5d**).

Additionally, we explored whether 3D electron microscopy patches sampled along neurite skeletons could be used directly for functional type prediction, bypassing morphological features (**Extended Data Fig. 5e**). We applied SegCLR (Dorkenwald *et al*., 2023), which trains a 3D ResNet-18 with contrastive loss (Chen *et al*., 2020) on patches extracted along neurite skeletons (16 nm/voxel, 49^3^), mean-pools patch embeddings per cell, and classifies with logistic regression. Using leave-one-out cross-validation on the FCLEM dataset (84 labeled cells, 4 functional types), SegCLR achieved a weighted F1-score of 32.9%. To test whether alternative patch representations and learning strategies could improve on this, we evaluated two further approaches: supervised contrastive learning (Khosla *et al*., 2021) on segmentation-masked cable patches at 8 nm resolution, which constrains the network to neurite-intrinsic ultrastructure (weighted F1 = 32.1%), and multiple instance learning with attention-weighted pooling (Ilse, Tomczak and Welling, 2018) on cable patches at 32 nm resolution, where the attention mechanism learns which patches along the neurite are most informative and produces a single cell-level embedding classified end-to-end with cross-entropy (weighted F1 = 33.5%). All three approaches performed substantially below the morphometric classifier (81.7%), suggesting that the limited number of labeled training cells is insufficient for learning discriminative representations directly from raw EM ultrastructure.

### Neuron reconstructions and neurotransmitter assignment in the WBEM dataset

To obtain high-quality, exhaustive reconstructions in the WBEM dataset, each neuron was reconstructed independently by at least two expert proofreaders using automatic segmentation algorithms in Neuroglancer. To minimize anatomical bias for selecting seed cells, we generated a list of IDs containing all automatically extracted cell bodies in the integrator region of the anterior hindbrain (containing rhombomeres 1 to 3) and randomly shuffled this list. We then attempted to completely reconstruct the axons and dendrites of each neuron in the shuffled list in sequence. Only neurons whose axons and dendrites terminated in natural endings within the volume (i.e., in arborizations or synaptic boutons) were deemed to be complete and were included as seed cells. Whenever tracing of axons or dendrites could not be continued due to issues with tissue quality, a neuron was discarded as a seed cell. After we had reached the predetermined number of 20 seed cells, we turned to the synaptic partner reconstructions. This very stringent quality criterion was slightly relaxed for the synaptic partner reconstructions because we had to optimize the level of completeness of reconstructions with the number of neurons that could be included in the dataset. Here, all neurons were included that could be reconstructed by at least two proofreaders independently to a ‘completeness’ of ~80%, when compared with the seed cells. This level of completeness proved to be sufficient for excellent LDA classification results. Neurons that passed these criteria were then added to the database of reconstructed morphologies and assigned a functional type by the LDA classifier, as described above. Neurotransmitter assignment is described in (Petkova, 2025).

### Synapse detection and synaptic partner reconstructions in the WBEM dataset

At the onset of the project, automatic synapse detection was not yet available in the WBEM dataset. Therefore, we opted for a completely manual synapse detection approach. Because manual synapse detection is generally considered to be ground-truth data and is used to benchmark automated synapse detection algorithms, we opted to continue using manual synapse detection even after automated detection became available. As with the neuron reconstructions, the dendrites and axons of every iMI seed cell were searched by at least two expert proofreaders for synaptic specializations, and a consensus list of synapses was generated. Then, we attempted to reconstruct the synaptic partner of the synapse of interest to its parent cell body. Our success rate for reconstructing dendrites of postsynaptic neurons was 72%, while the success rate for tracing the axons of presynaptic partners back to a parent cell body was 24%.

### Connectome-constrained recurrent neural network modeling

We modeled the bilateral circuit as a continuous-time rate recurrent neural network (RNN) with total units, implemented in PyTorch (torch v.2.9.0) and trained end-to-end via backpropagation through time. The network was organized into two hemispheres, each with neurons assigned to one of the four functional populations: iMI, cMI, MON, SMI. Each reconstructed hindbrain neuron whose functional identity was determined through direct functional imaging or LDA-based prediction was represented by a single unit in the network. Myelinated neurons present in the anatomical reconstruction (**Fig. 2a**) were excluded from the network architecture. Reconstructed rostral axons were used to define the input vector of the network. This one-to-one mapping determined the number of model units, their hemispheric and functional identities, and the connectivity among them and with external input, yielding a network of N = 182 neurons (**Fig. 5a**). We ran all model optimizations in parallel on the Baden-Württemberg high-performance compute cluster (bwForCluster NEMO 2) with 1 requested CPU core and 32GB RAM per model. Model optimization took approximately 2 hours on average, terminating when the loss stopped improving or the maximum of 5001 epochs was reached.

#### Forward simulation with exact exponential leak

The hidden-state dynamics, representing the membrane potential *h(t)* for each neuron in the network, can be understood as a discretization of the continuous-time leaky rate equation:

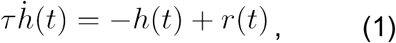

where *r(t)* is the “drive” component to the update, e.g. *r(t) =Wf(h(t))+Ux(t)*, combining the recurrent and input contributions. Given that the membrane state relaxes toward the instantaneous drive with a time constant τ (set to 0.1 s for all the nodes in the network and unaffected by training) and *dt* (set to 0.01 s for all the reported simulations in **Fig. 5** and **Extended Data Fig. 8**) is the discrete time step, the most direct explicit discretization of this equation is the forward Euler scheme:

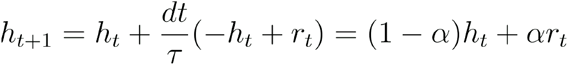

With 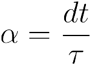, so the leak multiplier is 1 − α. In our implementation, however, we used the following equivalent exponential-form update:

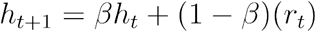

where

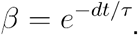

This can be derived by solving exactly **Eq. 1** over one time step while assuming that the total drive *r*_*t*_ remains constant during that step (Hochbruck and Ostermann, 2010). This procedure is often described as an exponential integrator or exact discretization of the linear part, and it is especially attractive for leaky systems because the decay associated with the membrane leak is represented exactly rather than approximated by a linear step. The link between the more traditional forward-Euler and the exponential form becomes clear by expanding the exponential factor for small *dt* : since *e*^*−dt/τ*^ *≈* 1 *− dt/τ* to first order, the exponential update reduces to the forward Euler rule in the limit of sufficiently fine time steps. Thus, both schemes approximate the same underlying differential equation, but the exponential version preserves the exact passive decay when the input drive is locally constant. Ultimately, the network state evolved according to the following leaky-integrator update rule with exact exponential discretisation:

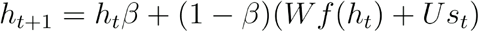

where *h*_*t*_ *∈ ℝ*^*N*^ is the vector of membrane potentials at time step t, *β=e* ^*−dt/τ*^ is the leak factor determined by the membrane time constant τ and simulation time step *dt*,*W ∈ ℝ*^*NxN*^ is the recurrent weight matrix, *U∈ ℝ*^*N*^ is the input weight vector, and *s*_*t*_ *∈ ℝ*^*N*^ is the external input at time. The firing rate of each unit was obtained by passing the membrane potential *h(t)* through a softplus nonlinearity

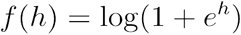

chosen for its non-negative output and smooth gradient.

#### Connectivity structure and biological constraints

We took a data-driven approach to the design of the network architecture. The number of neurons, functional and anatomical identity, excitatory or inhibitory (E/I) nature, and wiring diagram were all directly mapped from functional and connectomics information. The E/I labels were encoded in a 1-dimensional sign vector

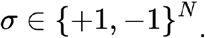

The recurrent connectivity weight matrix was then constructed as:

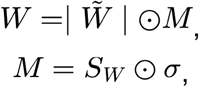

where the mask *M* encodes the biological constraints (**Fig. 5b**). The support matrix *S*_*W*_ *∈*{0.1}^*NxN*^ is the mask of zeros and ones, where ones describe the experimentally found connections.

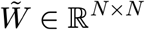 contains learnable raw weight parameters (initialised by drawing from 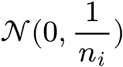 for neuron*i*, with *n*_*i*_ active input edges in *M*). ∣ · ∣ denotes elementwise absolute value to enforce non-negative weight magnitudes. The outer product, the elementwise multiplication by *σ* (denoted as ⊙) ensures that all outgoing weights from each unit share the same sign, in accordance with Dale’s law.

The input weight vector was constructed as

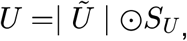

where Ũ is an array of learnable parameters, initialised drawing from 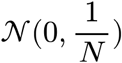. The support mask *S*_*U*_ *∈*{0.1}^*NxN*^ is defined by setting to non-zero only connections to neurons linked to at least one rostral axon in the connectomics dataset. Because the E/I identity of the corresponding presynaptic neurons is unknown, all input weights are constrained to be positive. This assumption prevents training from overriding the experimentally determined E/I character of neural interactions. For each population *p* with *N*_*p*_ neurons, the readout vector was defined as 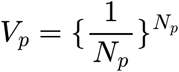. Concatenating *Vp* for all *p* produces the overall readout vector.

#### Population readout and GCaMP convolution

The network produced eight population output signals *y*^*p*^*(t)*, corresponding to the mean firing rate across all units in population *p* in a given hemisphere. To match the temporal filtering inherent in GCaMP-based calcium imaging, the raw firing rate traces *f*(*h*_*t*_) and population outputs *y*^*p*^*(t)* were convolved with a double-exponential GCaMP kernel characterised by a rise time constant *τ*_*rise*_=0.25*s* and a decay time constant *τ*_*decay*_ =2.4*s* prior to loss computation and output storage (Migault *et al*., 2018; Slangewal *et al*., 2026).

### Manipulations of the network structure for control tests

All manipulations of the network structure are performed before training. For each condition, 300 manipulated networks were produced. The distribution of their post-training loss was then compared using a two-sided Mann-Whitney U test against the loss distribution obtained from training the baseline network structure 300 times.

#### Shuffled connectivity

We assessed the contribution of the specific recurrent connectivity patterns in our dataset to reproducing the dynamics observed in the recorded neural traces. To this end, we trained networks with shuffled recurrent connectivity matrices. We shuffled only *S*_*W*_, leaving all other components of *W* unchanged to preserve each neuron’s E/I and population identity (**Extended Data Fig. 8d**). As a byproduct of this operation on the sparse connectivity matrix, important couplings between *W* and *U* could be compromised. We therefore allowed a full support vector *S*_*U*_=1_*N*_, enabling training to recover any disrupted couplings. Note that this additional flexibility favors the shuffled controls, making the comparison conservative and strengthening the evidence for the contribution of the original connectivity patterns to model performance. The readout vector *V* was kept unchanged.

#### Connections ablation

To further clarify the importance of the measured connectivity patterns in our model, we performed tests involving both targeted (**Extended Data Fig. 8**) and random removal of connections (**Extended Data Fig. 8**). These operations were performed only on the recurrent connectivity matrix *W*, while *U* and were kept unchanged. Targeted ablations were implemented by setting to zero a specified number of randomly selected elements within the left and right *iMI ↔ iMI* blocks of *S*_*W*_. For random ablations, the same operation was performed across the entire matrix *S*_*W*_.

### Training of recurrent neural network weights

Networks were trained by minimising the mean squared error (MSE) between the convolved simulated population outputs and the target calcium traces, weighted and summed across all *P*=8 populations and all trials for *M* training trials of duration *T*:

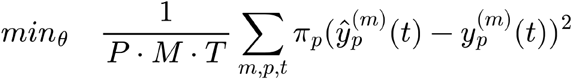

Where θ is the set of parameters including 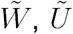. These were jointly optimised using the Adam optimiser with learning rate *η* =10^−3^ and weight decay *λ*=10^−5^. Gradients were clipped to a maximum norm of 1.0 to prevent instability. *π*_*p*_ are population weights designed to encourage the model to match the faster and slower dynamics of MON and SMI neurons respectively: *π*_*iMI*_ = *π*_*cMI*_ = 1, *π*_*MON*_ =2, and *π*_*sMI*_ =3.

#### Regularization

Two auxiliary penalty terms were added to the loss in a staged schedule:

1. *Spectral radius penalty*. A quadratic penalty was applied to the spectral radius *ρ* of *W* whenever it exceeded the limit *ρ*^*^=1 by a margin of 0.01: *ℒ*_*spec*_ = *λ*_*spec*_ [ReLU ( *ρ* (*W*) − *ρ*^*^ −0.01.)]^2^, encouraging the fast component to remain in a stable, globally sub-critical regime.
2. *Stimulus-gated antagonism penalty*. A penalty was applied to discourage configurations where the gain in the stimulated hemisphere was weaker than that of the contralateral hemisphere. This nudges the system towards an activity regime where lateralised slow dynamics are consistent with the stimulation side:

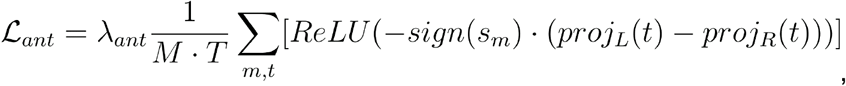

where *proj*_*L/R*_ *(t)* is the projection of the firing rate vector onto the summed left/right readout vectors extracted from *V*, and *sign* (*s*_*m*_) *∈*{+1−2} encodes stimulus laterality for trial *m*.

#### Training stages

Training proceeded in three stages over *N*_*epochs*_=5001 total epochs. In Stage 1 (first 30% of epochs), only a weak spectral penalty was applied, and the antagonism penalty was disabled (*λ*_*spec*_ =10^−3^,*λ*_*ant*_=0), allowing the network to first find a rough solution. In Stage 2 (next 30%), the full spectral penalty was engaged (*λ*_*spec*_ =10^−2^,*λ*_*ant*_=0). In Stage 3 (final 40%), the full antagonism penalty was additionally activated to shape hemispheric lateralisation ( *λ*_*spec*_ =10^−2^,*λ*_*ant*_=10^−5^). The gradual introduction of penalty terms alongside the core loss function made the training robust. The training trajectory is primarily driven by the optimization of the objective in the first stages. Once a suitable parameter subspace has been found, regularization nudges the network away from instability and from solutions with a single dominant side.

#### Training data

The model was trained jointly on the activity traces associated with *M*=4 stimulation conditions for each neuronal population (iMI, cMI, MON and SMI). The only experimentally acquired condition corresponds to leftward motion with a coherence of 100%. Here, each signal trace was obtained as the average *ΔF/F* signal across all recorded neurons belonging to the associated population. These traces were shifted to be positive throughout the entirety of the recorded window. This allowed each neuron’s firing rate to be computed as the output of a softplus function, which is strictly non-negative. The sign of each interaction was therefore determined solely by the corresponding connection weight, making its excitatory or inhibitory nature unambiguous. We augmented the training dataset by adding three more conditions: rightward motion at 100% coherence was obtained by mirroring the signals in the two hemispheres; left/rightward motion stimuli at 10% coherence (having 0.1 times the magnitude of the original stimulus signal) were obtained by scaling the original/mirrored response traces by 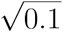 (with the sublinear relation accounting for Weber’s Law).

In order to avoid the noise contribution in the target signals being overrepresented, which could lead to overfitting, we computed a model of noise for the cMI populations and applied it to all raw traces corresponding to the three additional conditions. To define this noise contribution, we modeled the recorded activity trace within the time window of stimulation as the sum of a deterministic integrator component and a temporally correlated stochastic component. To this aim, we first isolated the deterministic component. This was modeled as a saturating exponential 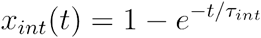, and the integration time constant *τ*_*int*_ was estimated by nonlinear least-squares fitting of this function to the windowed trace using SciPy v1.17.0 *curve_fit*. We then subtracted the fitted deterministic component from the data to obtain a noise-only signal and numerically differentiated it to approximate *x*. Assuming this residual follows a leaky integrator driven by additive noise, 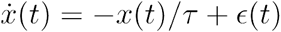, we estimated the leaky time constant *τ* by minimizing the Gaussian negative log-likelihood of the residuals with respect to *τ* using a bounded scalar optimizer (SciPy v1.17.0 *minimize_scalar*, bounded in the interval [10^−3^,100]*s*).

Here, *τ* determines the autocorrelation of the residual noise contribution. Instead of treating the resulting residual innovations *ε(t)* as the result of a Wiener process, they were more generally modeled as a discrete-time Ornstein–Uhlenbeck process, *ϵ*_*t+Δ t*_ = *α ϵ*_*t*_ +*η*_*t*_, with 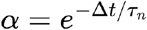 and *η ~ N(*0, *σ*^2^(1 *− α*^2^)). We estimated the OU timescale *τ*_*n*_ and noise amplitude *σ* by minimizing the exact Gaussian negative log-likelihood of the one-step transitions using a multivariate optimizer (SciPy v1.17.0 *minimize*). Using these fitted parameters, we simulated OU noise realizations and added them to the original experimental trace to generate signals with variable noise realizations.

#### Assessing single-neuron activity

The networks were trained to reproduce population-average activity rather than individual neuronal activity. So, we assessed whether individual neurons exhibited temporal profiles consistent with their assigned functional populations. For the training stimulus with 100% leftward coherence, we convolved the activity of single neurons with the GCaMP kernel, then computed Pearson correlations between each neuron’s activity trace and the corresponding population-average target trace. For each functional population, we computed the distribution of correlation coefficients across neurons from the top 2% of models ranked by loss post-training (**Extended Data Fig. 8c**). Then, we focused the analysis on the neurons in our models that contributed most strongly to shaping the population average activity. To this aim, we included in the distributions of correlation coefficients only neurons whose activity range (minimum to maximum activity amplitude of each neuron) fell within the highest quartile for their respective population.

## Data availability

All neurons described in this paper are browsable in FishExplorer: https://fishexplorer.zib.de/sandbox/. We have generated a website linked to the paper https://jboulanger91.github.io/fish1.5-release/, which provides access to interactive Neuroglancer visualizations of the EM dataset and Jupyter notebooks for retrieving neuronal morphology and connectivity.

Data related to this manuscript has been made available on the publicly accessible repository Zenodo under the persistent identifier https://zenodo.org/records/19231045. X-ray datasets are available through the Harvard Dataverse https://doi.org/10.7910/DVN/FFAACE.

Upon publication, source code and analysis scripts will be archived and made available on the publicly accessible repository Zenodo under a persistent identifier. During the review process, code is available on GitHub: https://github.com/jboulanger91/Zebrafish_CLEM.

## Author contributions

Project conceptualization: J.B.W., A.B.; Project lead: J.W.L., F.E., F.D.B., A.B.; EM Protocol development for FCLEM and WBEM datasets: J.B.W., M.P., R.L.S.; FCLEM sample preparation: J.B.W.; FCLEM sample imaging: J.B.W., R.L.S.; Development of EM image quality assessment tools: Y.W.; Electroporations: S.R.; Area photoactivations: S.R., A.B.; X-ray imaging and analysis: J.B.W.; FCLEM sample sectioning: R.L.S.; EM images stitching and coarse elastic alignment: Y.W.; Two-photon imaging and analysis: J.B.W., A.B., F.K., K.S., K.N.K.; 2P to EM image registration: J.B.W., A.B.; Neuronal tracing in FCLEM dataset: J.B.W., J.H.S.; FCLEM dataset connectome analysis: J.B.W.; Single-cell photoactivations: F.K.; Datasets synchronization: F.K.; Development and testing of LDA-based classifier: F.K., J.B.W., G.F.P.S., S.K.V.; Cross-modality prediction of functional morphotypes: F.K.; Rendering of neuronal morphologies: J.B.W., G.F.P.S., F.K., K.J.H.; Cross-modality neuronal morphology registration: A.B.; Modeling: R.G., A.B., A.L.F.; HCR FISH: F.K., H.N.; atlas management: S.K.V., D.B.; NBLAST quantification: M.E., F.K., S.K.V.; WBEM sample preparation: M.P., R.L.S.; WBEM sample imaging: M.P., J.B.W., R.L.S.; Neuronal tracing in WBEM dataset: G.F.P.S., K.J.H., R.T., M.S., A.H., D.H., Z.D., R.C.W., L.L.Z.; WBEM dataset connectome analysis: G.F.P.S., R.T., K.J.H., J.B.W., K.S.; H2B-GCaMP7f transgenic line generation: I.H.B.; cyto-GCaMP8s transgenic line generation: H.N.; Funding acquisition: J.W.L., F.E., F.D.B., D.B., A.B.; Conceptualization of figures: J.B.W., F.K., G.F.P.S., A.B., R.G.; Writing – original draft: All authors provided input to the writing of the manuscript; Writing – review: J.B.W., A.B., R.G., A.L.F.; Correspondence and requests for materials should be addressed to J.B.W. and A.B.

## Acknowledgments

We thank Zetta AI for alignment, cell segmentation, synapse detection & assignment, and hosting CAVE. We are grateful to Thomas Macrina for his assistance with connectome analysis, Paul Teufel for his initial help with segmentation, Zachary Miller for his contributions to manual registration, and Hillary Jean-Gilles for her help with tracing.

## Extended Data

**Extended Data Figure 1.**
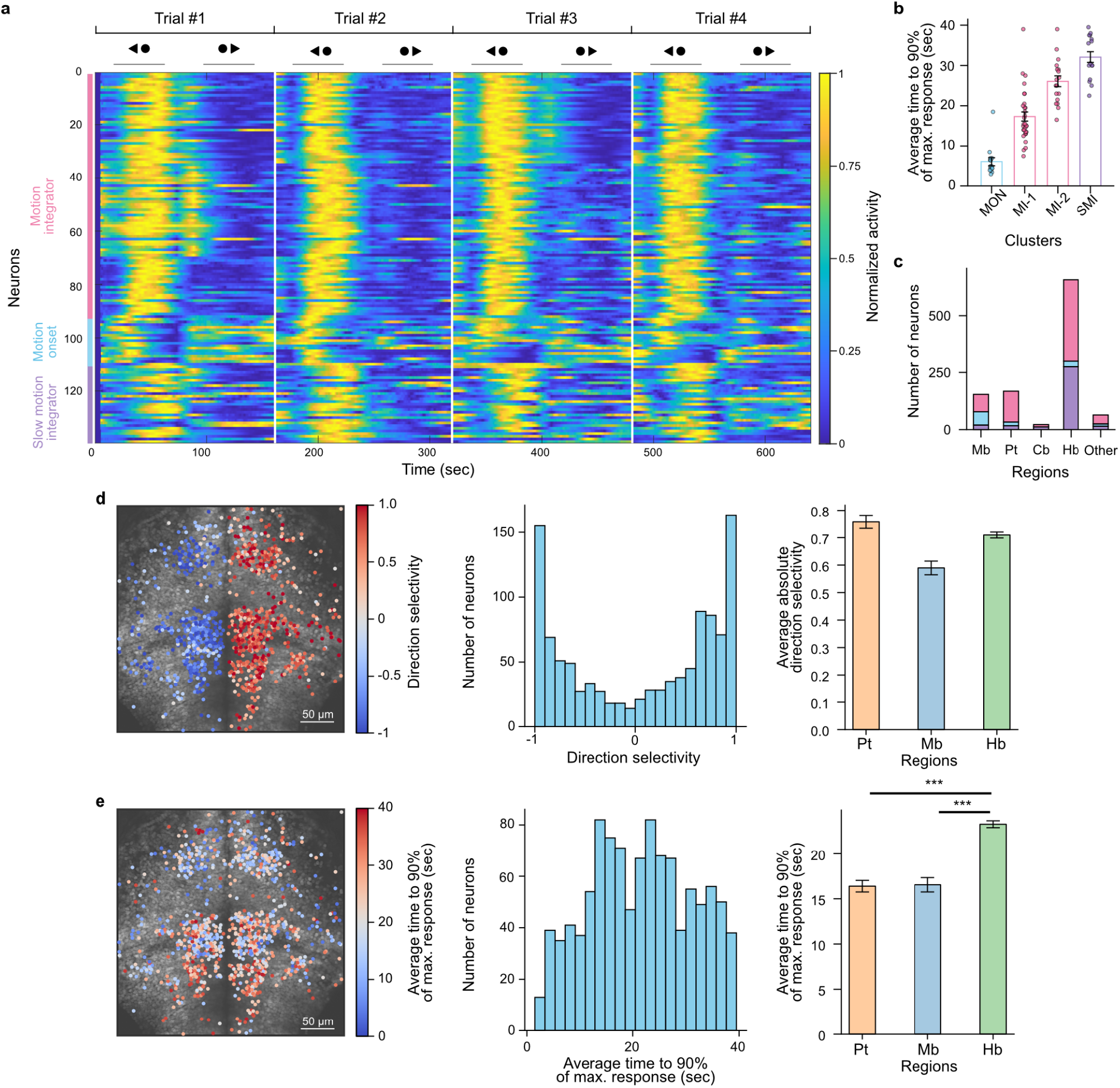
Detailed analysis of neuronal functional recordings. **(a)** Raster plot of representative examples of direction-selective neurons belonging to the three functional classes. Four example trials, both for left- and rightward-moving dots. The upper arrows represent the direction of motion, and the gray bars represent the stimulation epochs. Neurons are organized according to their functional responses (see left colored bars). Individual trials are separated by horizontal white bars. **(b)** Rise time to reach 90% of the maximal response for each k-means cluster. Each dot represents a single neuron. MON, motion onset neurons; MI-1, fast motion integrators; MI-2, intermediate motion integrators; SMI, slow motion integrators. **(c)** Distribution of neurons across brain regions and cluster types. (**d**) Left: spatial distribution of direction selectivity values for the three functional classes. Middle: histogram of direction selectivity values for the same population. Right: average values for neurons grouped by brain region. (**e**) Same as in (**d**), but showing the time to reach 90% of the maximal response., *** indicates p < 0.001 (two-sided t-test). Mb, midbrain; Pt, pretectum; Cb, cerebellum; Hb, hindbrain; Oth, other regions.

**Extended Data Figure 2.**
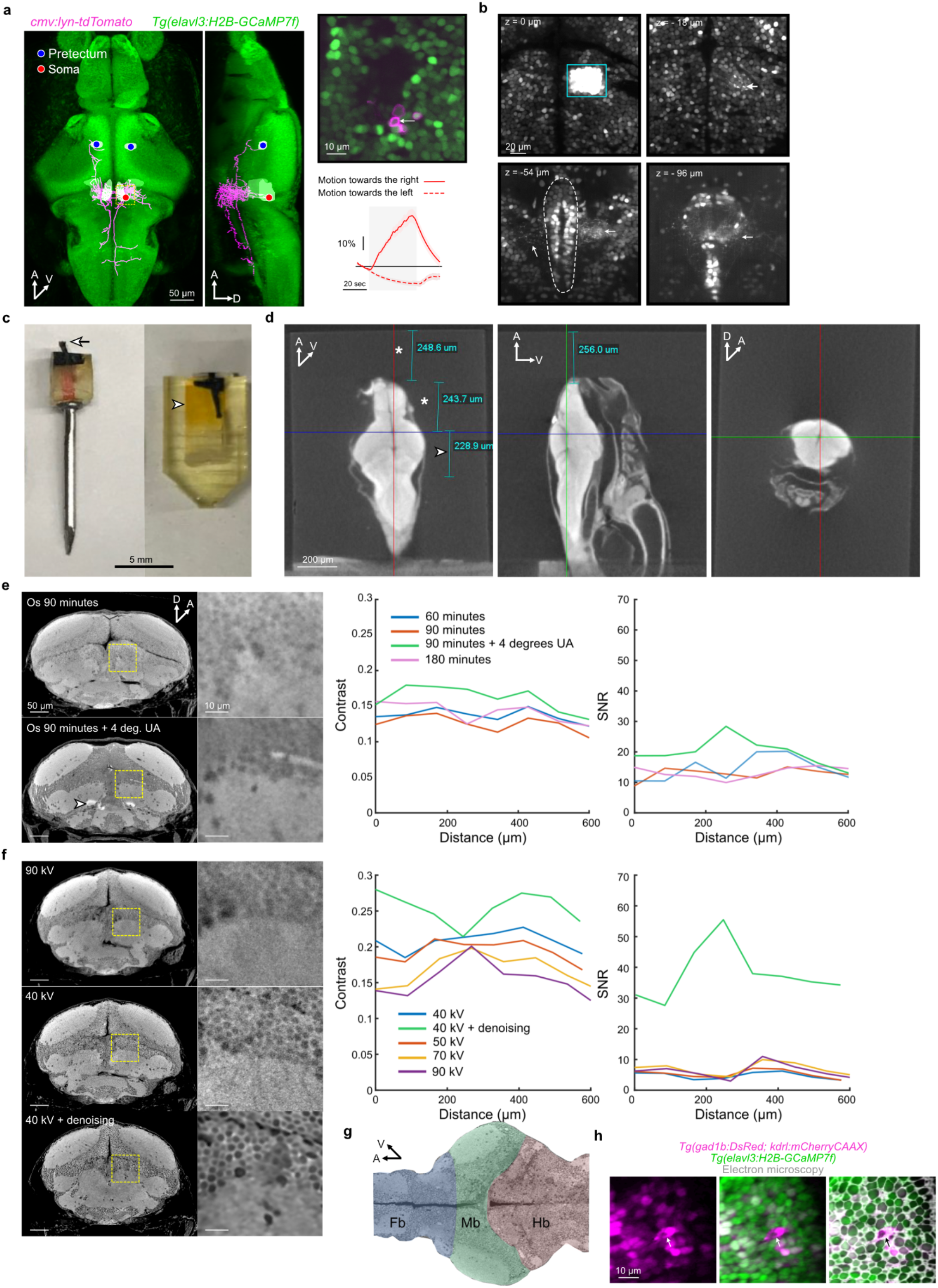
Sample preparation and region of interest identification. **a**, hindbrain sparse electroporations. Left panels: reconstructed neurons registered to a reference brain, indicating soma and Pt locations. Note the projection pattern of the cells towards the Pt and the ipsilateral and contralateral anterior hindbrain. The dashed yellow square is magnified in the top-right snippet. Right panels: two-photon single-plane image showing the somas of the electroporated neurons together with response dynamics to rightward and leftward moving dots for one neuron (indicated by the white arrow). The activity responses are reminiscent of an evidence integrator. Activity traces show the average responses for 4 trials (SEM indicated by the red-shaded area). The stimulation epoch is denoted by the shaded gray area. The observed pre-stimulus decrease reflects fluorescence decay from the preceding stimulation trials, as trials were presented in close temporal succession to increase statistical power. **b**, Photoactivation of a group of neurons in the anterior hindbrain in *Tg(elavl3:H2B-GCaMP6s, gad1b:loxP-DsRed-loxP-GFP, alpha-tub:c3pa-GFP)* (cyan square). Cells project ventrally, ipsilaterally, and contralaterally close to the raphe (dashed outline at z=–54 µm from the photoactivation plane). Neuronal projections are indicated by the white arrows. **c**, Left: A resin-embedded larva with the cast around the head trimmed (white arrow) for enhanced X-ray penetration. Right: Another larva re-embedded in resin after X-ray tomography before ultramicrotome sectioning. The limit between the two resin casts is denoted by the arrowhead. **d**, Orthogonal X-ray tomography views of the main larva used in this study. The first panel shows the edges of the region of interest determined before the section collection. By performing direct measurements using the X-ray microscope software, we precisely targeted the section collection by trimming away the unnecessary resin and tissue (white asterisks), keeping only the region of interest (white arrowhead) containing the anterior hindbrain and most of the identified neuronal projections (**a,b**). **e**, Optimization of the EM staining protocol using independent specimens that were not subsequently sectioned for EM. X-ray tomography was used to assess improvements in image contrast and signal-to-noise ratio following changes in osmium staining duration and uranyl acetate incubation conditions. The first step of contrast and signal-to-noise ratio (SNR) improvement was achieved by varying the Osmium (Os) incubation duration and incubating the Uranyl Acetate (UA) at 4°C instead of room temperature. The arrowhead indicates staining artefacts. Left panels: Representative X-ray tomography cross sections along the main axis of the larva and high-resolution snippets (corresponding to dashed yellow squares). Right panels: Contrast and SNR computed along the main axis of the fish, see **Extended Data Figure. 2g**. Each line represents the average of two fish per condition. **f**, X-ray imaging of the separate specimen subsequently used for FCLEM, showing improved image quality after optimization of the X-ray imaging power. Notice the dose-dependent improvement as the voltage is lowered from 90 to 40 kV. X-ray volumes corresponding to **e** and **f** can be found here. **g**, Reference fish imaged with X-ray (dorsal section plane) indicating where the contrast and SNR measurements were performed. Distances correspond to the upper plots. Fb, forebrain; Mb, midbrain; Hb, hindbrain. **h**, Images showing the alignment between the 2P and EM datasets confirming the precise correspondence of a blood vessel cross-section (indicated by a white arrow) across imaging modalities. Abbreviations: A, anterior; D, dorsal; V, ventral.

**Extended Data Figure 3.**
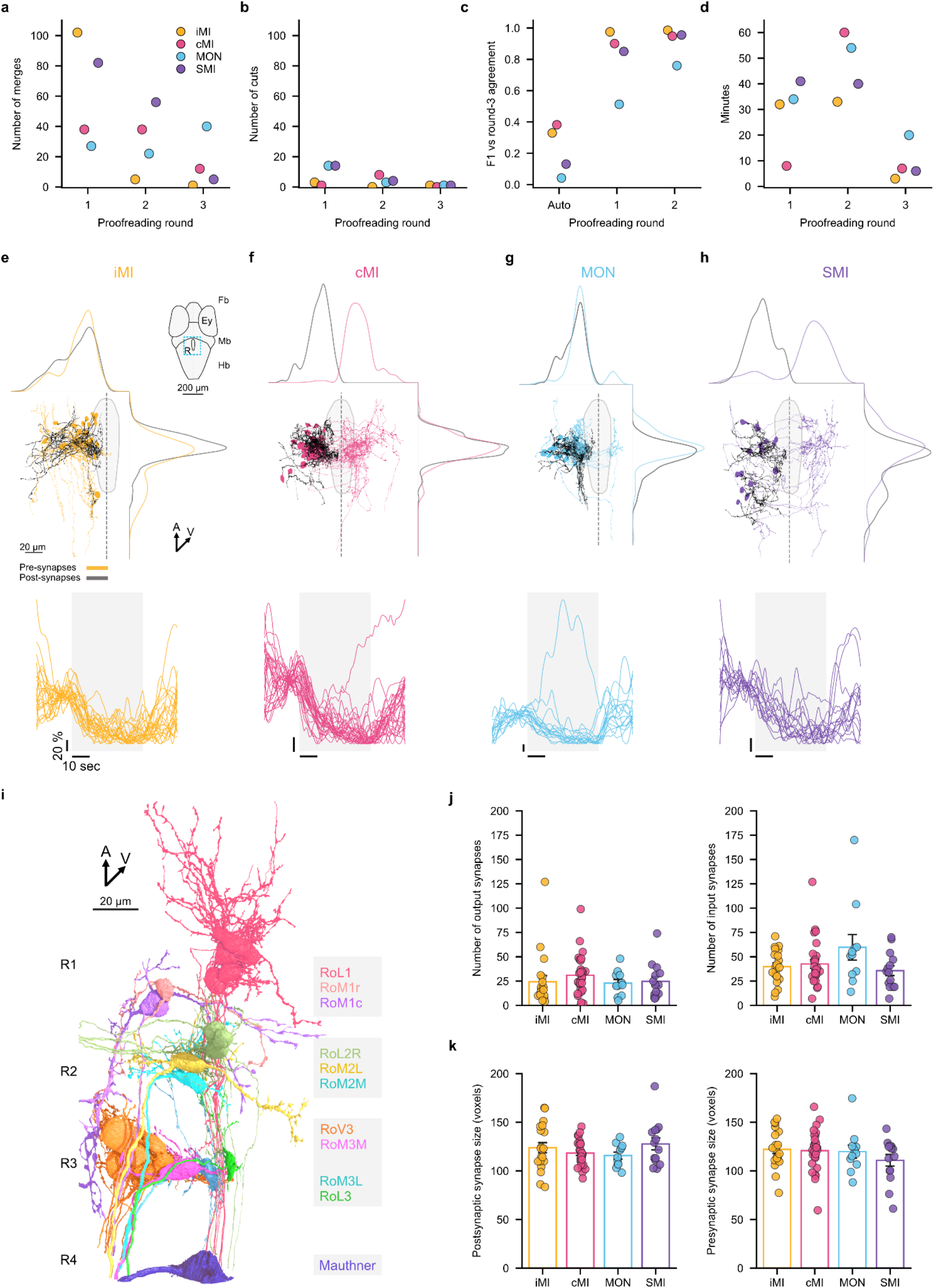
Supporting information for reconstructed neurons. **a-d**, Proofreading edits, synapse retrieval, and tracing time for 4 example neurons. Each point represents one neuron, colored by functional class: iMI, cMI, MON, or SMI. The first two panels show the number of merges and cuts recorded during proofreading rounds 1–3. The third panel shows synapse-retrieval F1 scores for the automatically generated base segmentation (“Auto”) and rounds 1–2, calculated against the round-3 agreement state as ground truth. The fourth panel shows tracing time in minutes for rounds 1–3. **e**, Structure-function relationship of functionally imaged and reconstructed iMI neurons as in **Fig. 1f–i** but in dorsal view. Somas and axons are colored in orange and dendrites in black. The dashed line represents the midline. Surrounding plots are the distribution of presynaptic (orange) and postsynaptic (black) synapses along the x (top) and y (right) axes. Right: schematic showing coarse brain organization (dorsal-to-ventral view) and location of the reconstructed neurons below (blue dashed square), surrounding outlines of the raphe (also shown below). Below: traces representing the normalized ΔF/F_0_ neuronal activity over time for motion in the opposite, null direction. **f–h**, identical representations for other neuron types. All neurons are registered to a reference brain (Randlett *et al*., 2015). **i**, Rendering of identified reticulospinal neurons belonging to rhombomeres 1 to 4 (R1–4). **j**, Bar plots showing the distributions of synapse numbers and sizes (**k**) for each functionally identified cell type. Error bars represent the standard error of the mean (SEM). Individual dots represent single neurons. Abbreviations: Mb, midbrain; Hb, hindbrain; R, raphe; Ey, eye; A, anterior; V, ventral.

**Extended Data Figure 4.**
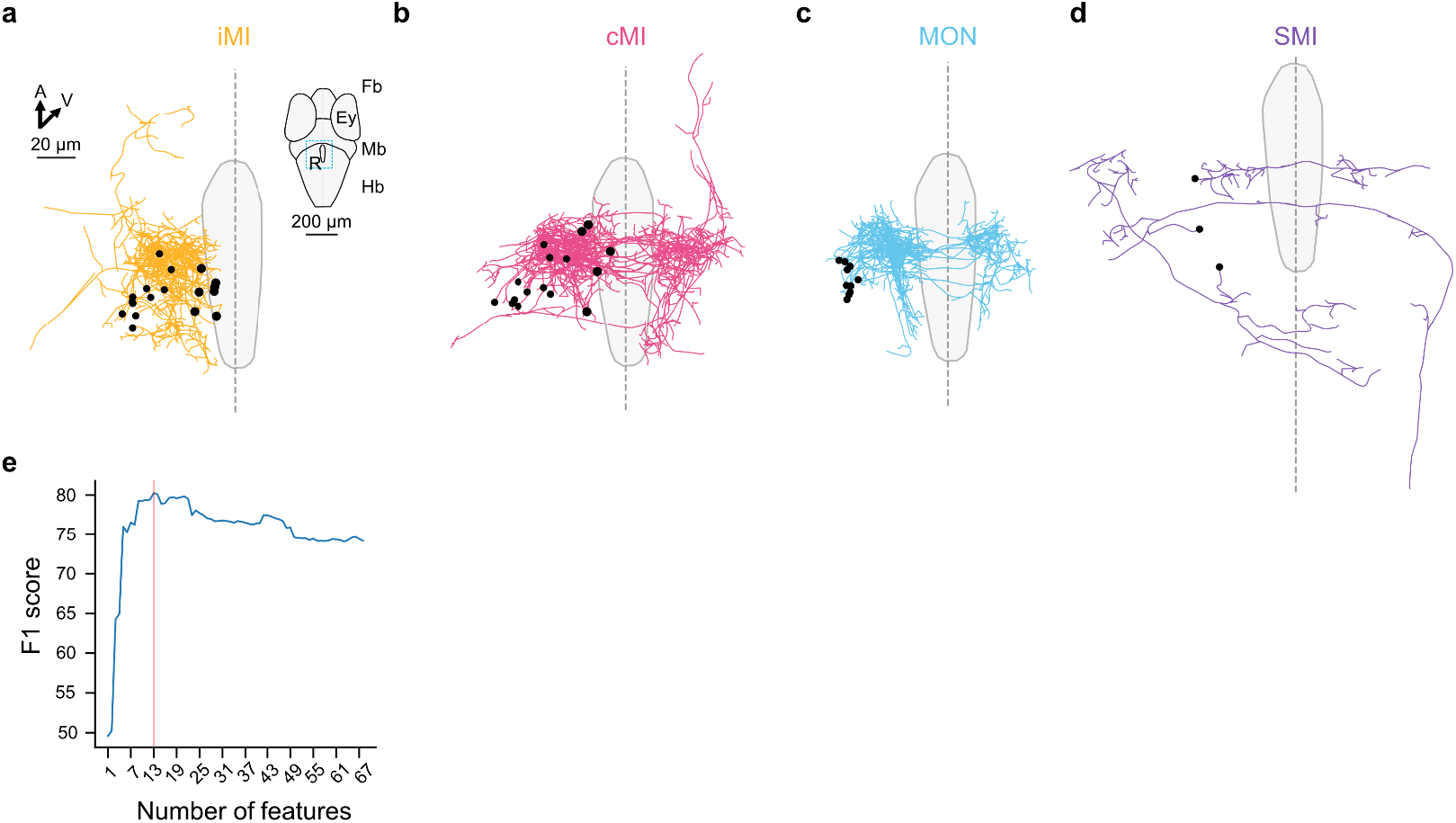
Diversity of photoactivated neurons and number-of-features selection. **a**, Structure-function relationship of functionally imaged and photoactivated neurons as in **Fig. 3b–e** but in dorsal view. Right: schematic showing coarse brain organization (dorsal-to-ventral view) and location of the reconstructed neurons below (blue dashed square) and outlines of the raphe (also shown below). **b–d**, identical representation for the other neuron types. All neurons are registered to a reference brain (Randlett *et al*., 2015). Abbreviations: Mb, midbrain; Hb, hindbrain; R, raphe; Ey, eye; A, anterior; V, ventral. **e**, F1-score as a function of the feature subset size used for **Fig. 3g** and below. The red line represents the number of features (n=13) yielding the maximum F1-score.

**Extended Data Figure 5.**
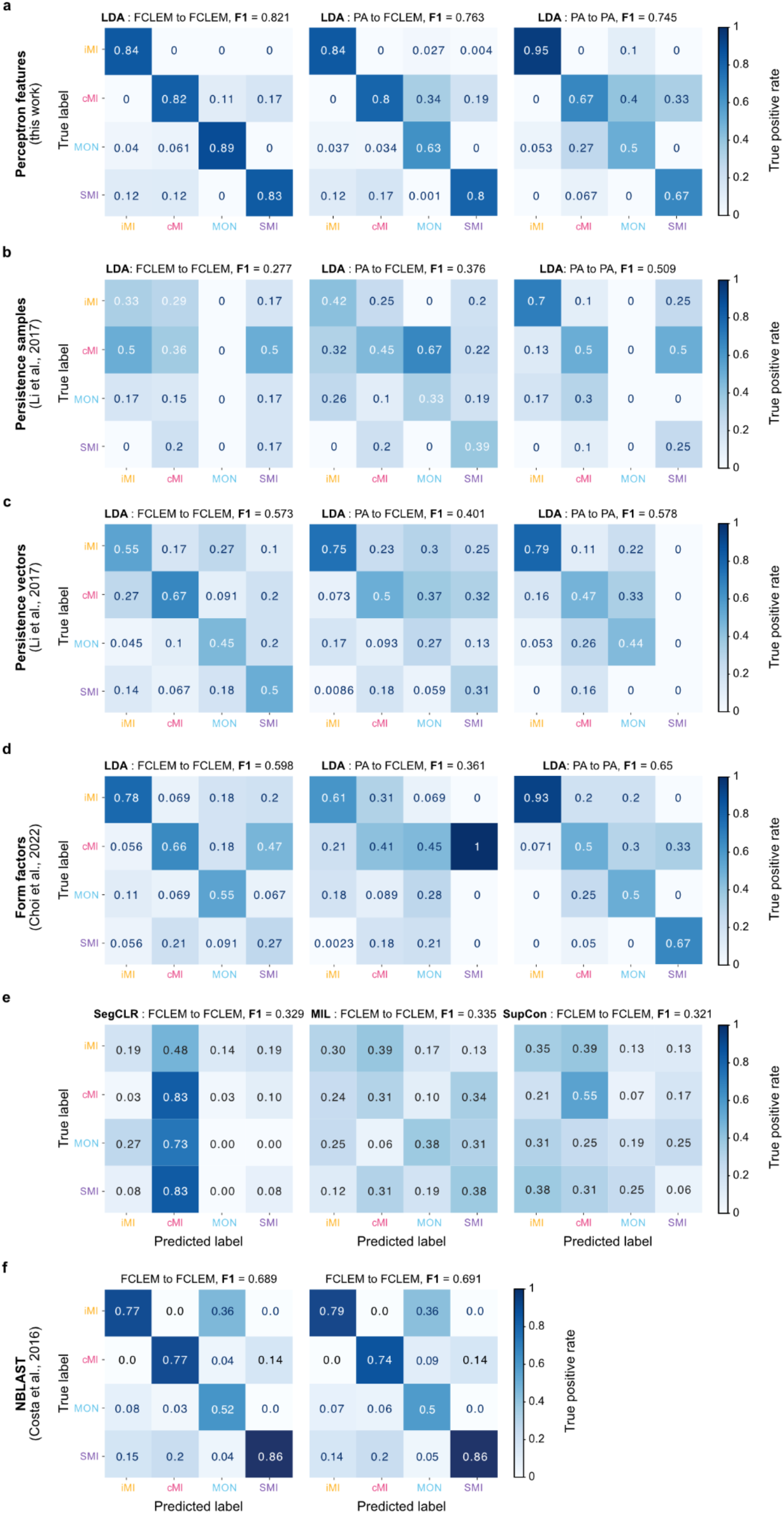
LDA benchmarking. **a**, Left, LDA confusion matrix of true positive rate using FCLEM cells as the ground truth and predicting FCLEM cells. F1-score for the entire matrix is indicated in the title. Middle, same as left, but using PA as ground truth and predicting FCLEM cells. Right, same as left, but using PA as ground truth and predicting PA cells. LDA, linear discriminant analysis; RF, random forest. **b–d**, Prediction of functional types using other established methods (persistence samples, persistence vectors, form factors metrics, respectively). In **e**, SegCLR corresponds to (Dorkenwald *et al*., 2023), MIL corresponds to multiple-instance learning with attention-weight pooling (Ilse, Tomczak and Welling, 2018), and SupCon to supervised contrastive learning (Khosla *et al*., 2021). **f**, Left, NBLAST confusion matrix generated using morphological similarity distances computed using the original *Drosophila*-based reference matrix (Costa *et al*., 2016) and right, the functionally identified zebrafish neurons in this study (**Methods**). Matrices were generated from a dendrogram using a cutoff required to separate 4 clusters.

**Extended Data Figure 6.**
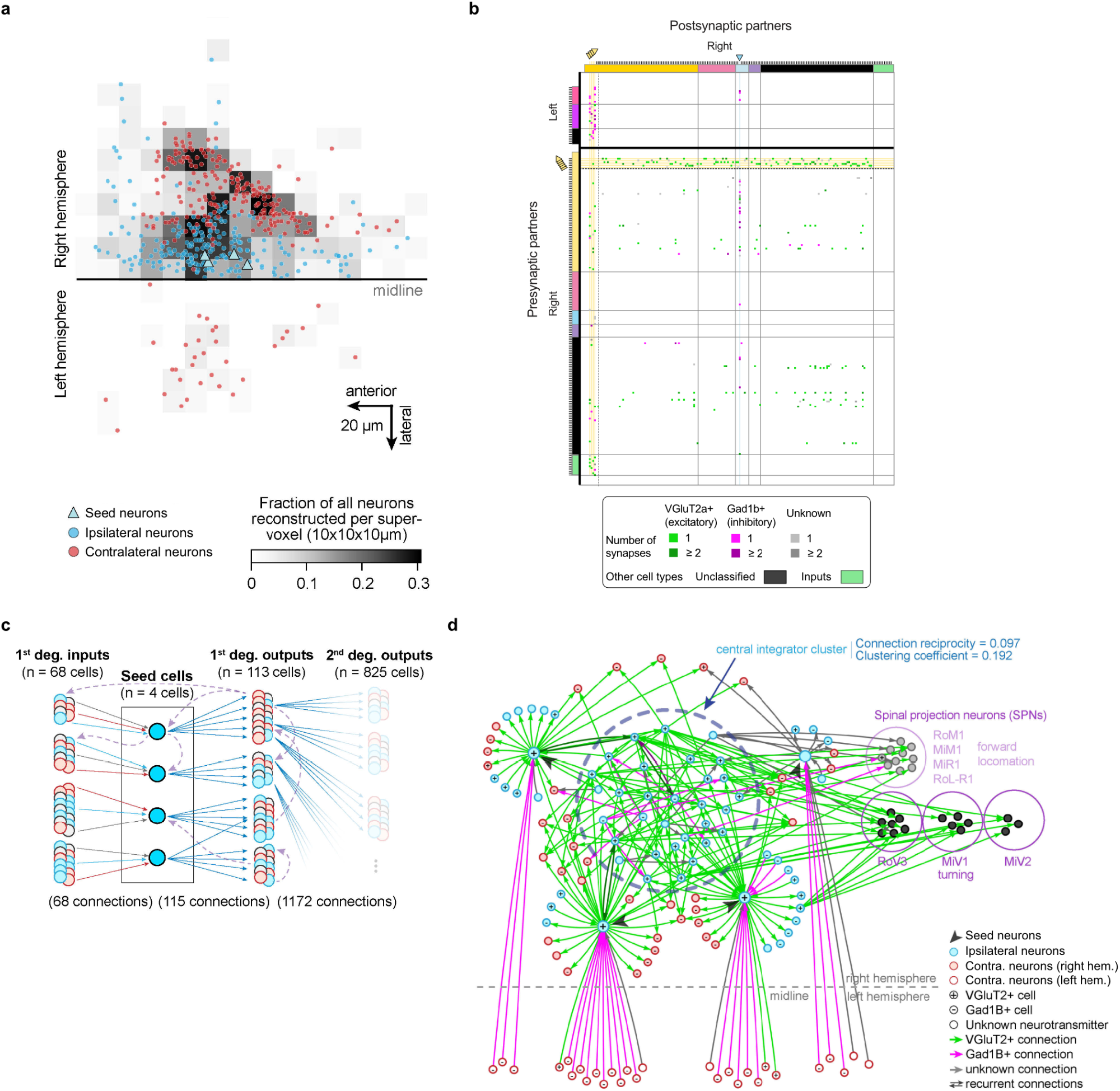
Supporting data for recurrent connectivity analysis. **a**, Z-projection displaying the reconstruction density (greyscale) achieved with our tracing approach in the anterior hindbrain; location of seed cells, z-projection contains 6 sections (i.e., 60 μm in Z). To obtain the reconstruction density, the anterior hindbrain was parcellated computationally into supervoxels (10 x 10 x 10 μm) and we divided the number of cell bodies we had reconstructed in each supervoxel by the total number of cell bodies located in this supervoxel. **b**, Connectivity matrix for LDA-classified WBEM neurons, organized by morphotype and hemisphere as in **Fig. 4g**, but also with unclassified neurons (black bars). **c**, Schematic depiction of our strategy for exhaustive circuit reconstruction of inputs and outputs of four iMI seed cells; blue, ipsilaterally-projecting cells; red, contralaterally-projecting neurons; grey, other cell classes; solid arrows, reconstructed synaptic connections (same color code); dashed arrows, recurrent ‘within-network’ connections revealed by circuit reconstruction. **d**, Network graph showing seed cells are connected with each other via a highly recurrent central integrator cluster. Connections are color-coded by neurotransmitter type.

**Extended Data Figure 7.**
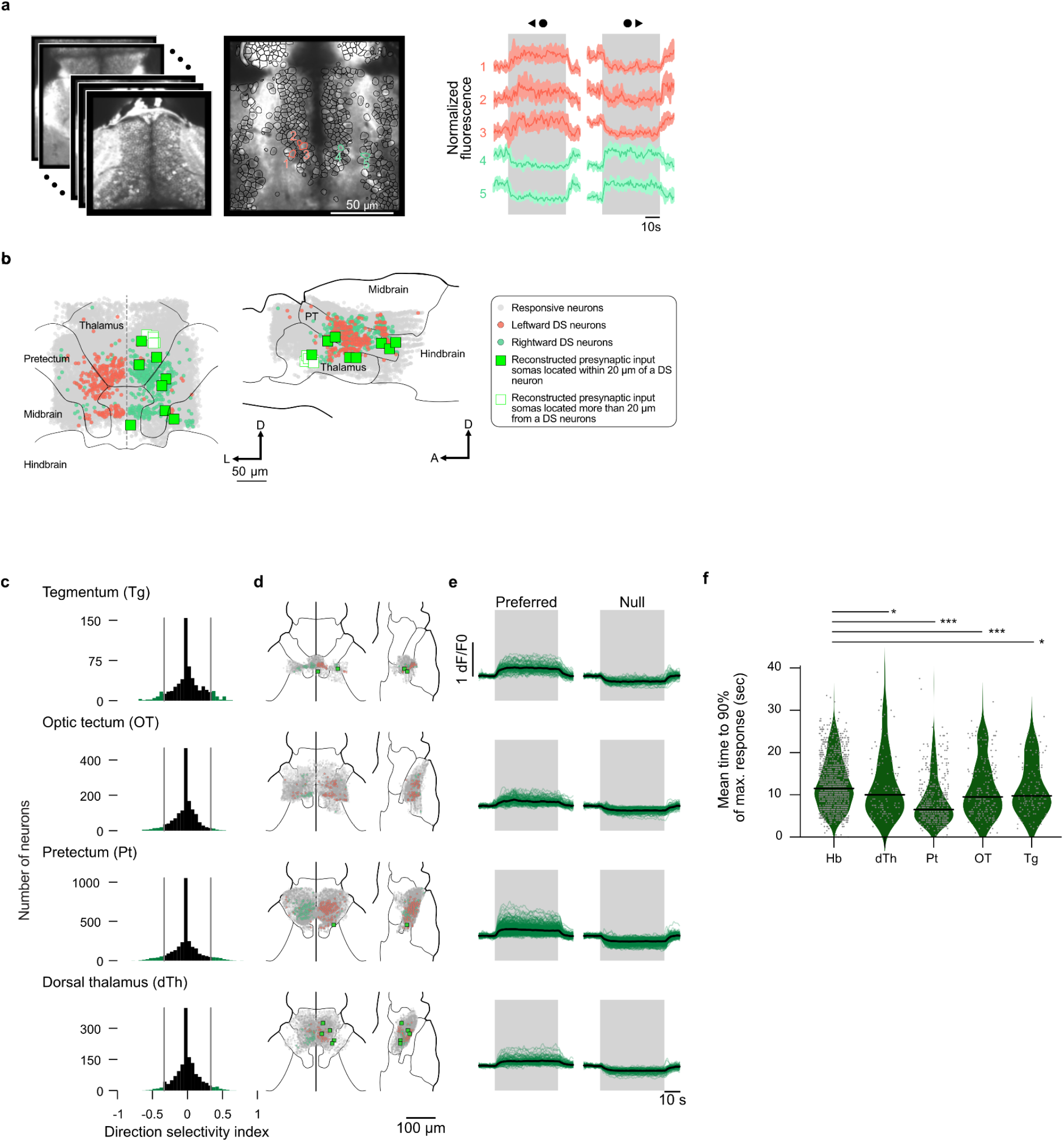
Imaging of thalamic, pretectal, and midbrain direction-selective inputs. **a**, Volumetric two-photon imaging (left) during random dot motion with example direction-selective units (right) for n=5 larvae. Horizontal gray bars represent stimulation epochs. Solid line and colored confidence intervals are the median and the interquartile range. **b**, Reference brain-registered functionally imaged and traced WBEM neurons, showing proximity of direction-selective units and iMI inputs (filled green squares). Annotated reference brain regions have been taken from the mapZebrain atlas ((Kunst *et al*., 2019), and **Methods**). **c**, Distribution of direction selectivity (DS) index for recorded neurons responding to coherently moving random dots in several regions anterior to the hindbrain. Histogram bins with DS superior to 0.33 or inferior to –0.33 are colored in green. **d**, Spatial distribution of neurons split by brain regions. Legend identical to that in **b. e**, functional responses in both the preferred and the null direction. Green lines indicate trial-averaged single neuron responses, and black lines indicate averages per brain region. **f**, Comparison of the rise time constant for the regions in **c–e** quantified as the mean time to 90% of the maximum response in seconds using one-sided Mann-Whitney U tests with Holm-Bonferroni correction. * alpha=0.05, ** alpha=0.01, *** alpha=0.001. Abbreviations: Pt, pretectum; L, lateral; D, dorsal; A, anterior.

**Extended Data Figure 8.**
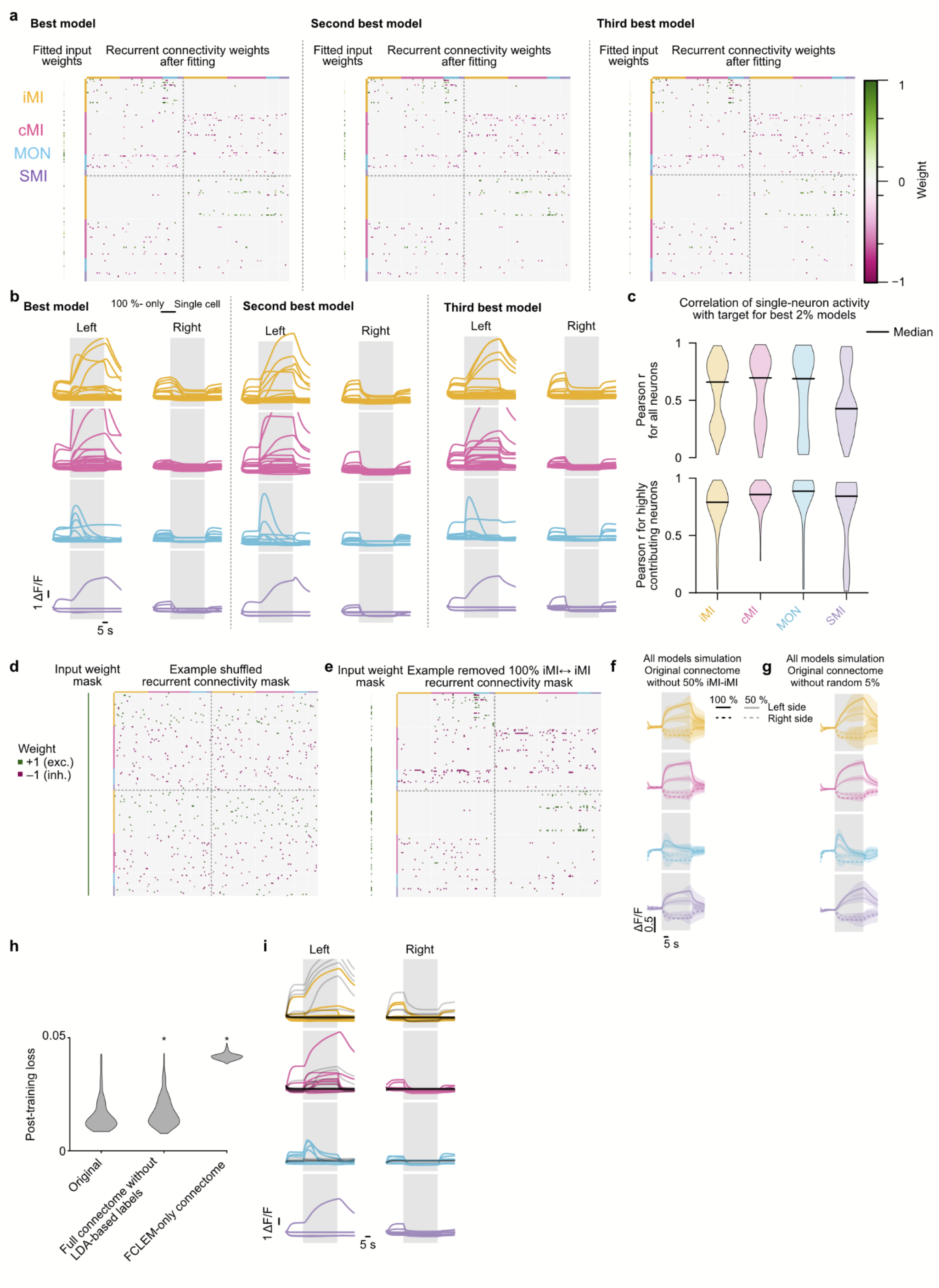
Details for connectome-constrained network fitting and model validation. **a**, The recurrent and input connectivity matrices of the optimized three best network models, following the connectome-constrained masking in **Fig. 5b. b**, Dynamics of all individual model neurons for each functional cell type on the left and right hemispheres in response to leftward coherent motion (gray box). **c**, Distributions of Pearson correlations between the target functions in **Fig. 5a** and individual neuronal activity traces from the top 2% of models, grouped by functional cell type. Correlation coefficient distributions are shown for all neurons (top) and for neurons with activity ranges in the highest quartile within each population, designated as substantially contributing to the population average (bottom). In iMI, these neurons showed a median correlation coefficient (black lines) r_m_=0.79 and an inter-quartile range r_IQR_=0.19; in cMI r_m_=0.86 and r_IQR_=0.12; for MON r_m_=0.89 and r_IQR_=0.11; finally for SMI r_m_ = 0.84 and r_IQR_= 0.23. **d**, Example shuffled connectome-constrained recurrent connectivity mask. Shuffling preserved neurotransmitter identities by initially mixing all connections, followed by cell-specific reassignment of excitatory and inhibitory labels. To make the comparison with the original connectome-derived models more conservative, all possible input connections were permitted for models with shuffled connectivity. This enabled training to restore couplings between input and recurrent connections that may have been disrupted by shuffling. **e**, Example recurrent connectivity mask with all iMI to iMI connections removed. Note that only synapses were removed, rather than entire cells in the network. **f**, Functional dynamics after removal of 50 % of iMI-to-iMI connections. Average ± std for all 300 optimized models. Median correlation coefficient between network readout and experimental traces across models and populations is r_m_=0.74, with r_IQR_=0.16. **g**, Same as in (f) but with 5 % randomly removed connections throughout the network. Note the smaller impact on slow integration dynamics in this case. Median correlation coefficient between network readout and experimental traces across models and populations is r_m_=0.90, with r_IQR_=0.02. **h**, Loss distributions after training for the original connectome-constrained model and two control variants. In control variant 1, we did not functionally label neurons with LDA but kept these unknown cells in the network during fitting (**Fig. 3f–h**). In control variant 2, we did not use these neurons, keeping only the original FCLEM dataset (**Fig. 2b**). Both variants lead to lower model fit quality (larger loss). Two-sided Mann-Whitney U test yielded p < 0.001, N = 300 fitting repeats each. **i**, Although neurons in control variant 1 (h) were excluded from the loss computation, integrator neurons that contributed substantially to the population activity exhibited dynamics consistent with their functional cell type. Non-labeled neurons in gray, FCLEM-based labels in color. Categorization of non-labeled neurons based on the LDA classifier.

**Extended Data Table 1.** All extracted 68 neuron morphometries used to generate our functional cell type classifier.

| # | Name | Extraction strategy |
| --- | --- | --- |
| 1 | number of contralateral branches | custom code |
| 2 | number of ipsilateral branches | custom code |
| 3 | **cable length** | navis |
| 4 | **bbox volume** | navis |
| 5 | **x_extent** | navis |
| 6 | **y_extent** | navis |
| 7 | z_extent | navis |
| 8 | **x average** | custom code |
| 9 | **y average** | custom code |
| 10 | z average | custom code |
| 11 | soma x | custom code |
| 12 | soma y | custom code |
| 13 | soma z | custom code |
| 14 | tortuosity | navis |
| 15 | n_leafs | navis |
| 16 | n_branches | navis |
| 17 | n_ends | navis |
| 18 | n_edges | navis |
| 19 | main_branchpoint | navis |
| 20 | n_persistence_points | navis |
| 21 | max_strahler_index | navis |
| 22 | sholl_distance_max_branches | navis |
| 23 | sholl_distance_max_branches_cable_length | navis |
| 24 | sholl_distance_max_branches_geosidic | navis |
| 25 | sholl_distance_max_branches_geosidic_cable_length | navis |
| 26 | main_path_longest_neurite | custom code |
| 27 | main_path_total_branch_length | custom code |
| 28 | first_major_branch_longest_neurite | custom code |
| 29 | first_major_branch_total_branch_length | custom code |
| 30 | first_branch_longest_neurite | custom code |
| 31 | first_branch_total_branch_length | custom code |
| 32 | cable_length_2_first_branch | navis |
| 33 | z_distance_first_2_first_branch | navis |
| 34 | biggest_branch_longest_neurite | custom code |
| 35 | biggest_branch_total_branch_length | custom code |
| 36 | longest_connected_path | custom code |
| 37 | n_nodes_ipsi_hemisphere | custom code |
| 38 | n_nodes_contra_hemisphere | custom code |
| 39 | n_nodes_ipsi_hemisphere_fraction | custom code |
| 40 | n_nodes_contra_hemisphere_fraction | custom code |
| 41 | **x_location_index** | custom code |
| 42 | **fraction_contra** | custom code |
| 43 | **y_extent_ipsi** | custom code |
| 44 | **z_extent_ipsi** | custom code |
| 45 | **max_x_ipsi** | custom code |
| 46 | **max_y_ipsi** | custom code |
| 47 | max_z_ipsi | custom code |
| 48 | min_x_ipsi | custom code |
| 49 | min_y_ipsi | custom code |
| 50 | min_z_ipsi | custom code |
| 51 | max_x_contra | custom code |
| 52 | max_y_contra | custom code |
| 53 | max_z_contra | custom code |
| 54 | min_x_contra | custom code |
| 55 | min_y_contra | custom code |
| 56 | min_z_contra | custom code |
| 57 | avg_delta_death_birth_persitence | navis |
| 58 | median_delta_death_birth_persitence | navis |
| 59 | std_delta_death_birth_persitence | navis |
| 60 | z_extent_contra | custom code |
| 61 | y_extent_contra | custom code |
| 62 | midline crossing x | custom code |
| 63 | midline crossing y | custom code |
| 64 | midline crossing z | custom code |
| 65 | crossing angle | custom code |
| 66 | crossing angle 2D | custom code |
| 67 | **morphology (ipsilateral or contralateral)** | custom code |
| 68 | neurotransmitter (exc., inh., or unknown) | custom code |

The list of all 68 morphometrics we extracted from each cell to build our functional cell type classifier. We used the navis package to compute features directly from .swc files and developed custom code to further extend feature space. Through a feature ranking with recursive feature elimination algorithm (see **Methods**), we identified 13 features to be the best predictor of function (highlighted in red).

